# Plumage diversity without differential expression of pigmentation genes in a rapid avian radiation

**DOI:** 10.64898/2026.11.05.764602

**Authors:** María Recuerda, Márcio Repenning, Matthew B. Toomey, Simón Kraemer, Jonas R. R. Rosoni, Luis F. Silveira, Cecilia Kopuchian, Leonardo Campagna

## Abstract

Uncovering the molecular basis of reproductive traits is key to understanding species divergence. Capuchino Seedeaters (*Sporophila*), a radiation of genetically similar birds differing mainly in male reproductive plumage color, can be leveraged to study how such traits evolve. Outlier scans identified candidate pigmentation variants primarily in noncoding sequence, implying regulatory changes. However, RNA-seq of growing feather follicles generally did not show differential expression of pigmentation genes among different colors, with only *ASIP* downregulated in crown feathers. Gene expression patterns instead varied by body region. The largest difference between dimorphic sexes was the upregulation of the carotenoid-degrading gene *BCO2* in males, accompanied by lower carotenoid content in male feathers. An improved genome annotation assigned 9.5% of candidate variants into coding and untranslated sequence. This suggests species-diagnostic variants lie within transcribed sequence and could act post-transcriptionally, rather than through transcript abundance, leading to a revised hypothesis for plumage divergence in this radiation.

**Summary:** Species-specific plumage patterns in males from different species belonging to the rapid Capuchino Seedeater radiation function as premating reproductive barriers. Genetic differentiation among species is concentrated in non-protein-coding regions near pigmentation genes, yet does not lead to the differential expression of these genes. Rather, species-diagnostic variants on the transcripts themselves, mostly in untranslated sequence, likely shape color phenotypes in the radiation.

## INTRODUCTION

Understanding the processes that generate phenotypic diversity is key to explaining how species diverge. Rapid radiations, in which closely related species diverge dramatically in morphology, coloration, or behavior despite minimal genome-wide differentiation, provide powerful natural experiments for identifying the molecular mechanisms underlying such phenotypic change (*1*, *2*). Growing evidence suggests that changes in gene regulation frequently underlie these rapid phenotypic transitions, often more so than protein-coding divergence (*3–6*). However, establishing links between genomic variation and regulatory evolution requires not only the identification of candidate loci, but also accurate genome annotations and transcriptomic data capturing gene expression in the relevant developmental stages and tissues (*7*).

Avian melanin-based plumage coloration provides a tractable system for investigating the molecular basis of phenotypic evolution because pigment production occurs within developing feather follicles, allowing gene expression to be measured directly during feather formation in adult individuals (*8*). Black, brown, and rufous plumage is determined by the balance between eumelanin and pheomelanin synthesis within follicular melanocytes (*9*). A central regulator of this process is the Agouti Signaling Protein (*ASIP*), which antagonizes *MC1R* signaling to promote pheomelanin production and has repeatedly been implicated in the evolution of pigmentation patterns across vertebrates (*10*).

The Capuchino Seedeaters (*Sporophila*) represent a striking example of rapid phenotypic diversification. This recent radiation comprises twelve closely related South American species that differ primarily in adult male plumage coloration and song, while exhibiting remarkably low genome-wide differentiation (*11–13*). Population genomic analyses have consistently shown that divergence among species is concentrated within a small number of narrow, highly differentiated genomic regions located near melanogenesis genes, including *ASIP*, *TYRP1*, *OCA2/HERC2*, *SLC45A2*, and *MLANA*, suggesting that regulatory evolution (rather than changes leading to protein differences) at these loci plays a major role in generating species-specific plumage patterns (*12*, *14*, *15*). Despite these findings, it is still unclear if such genomic differences lead to changes in gene expression among different feather patches and species, ultimately producing distinct color patterns. Directly testing this requires measuring gene expression in the developing feather tissues where pigment is produced.

A high-quality genome annotation is fundamental for interpreting both transcriptomic and population genomic data. Incomplete annotations can underestimate gene expression, miss alternatively spliced transcripts, and misclassify functional variants as intronic or intergenic, obscuring their biological function and significance (*16*). Recent advances in long-read transcript sequencing have substantially improved genome annotation by resolving complete transcript structures, alternative isoforms, and untranslated regions that are difficult to reconstruct from short-read RNA sequencing alone (*17*). Integrating Pacific Biosciences (PacBio) Iso-Seq with short-read RNA-seq has therefore become a powerful strategy for refining genome annotations and improving the functional interpretation of genomic variation, particularly in non-model organisms where reference annotations remain incomplete (*18–20*). Improving the annotation of the *Sporophila hypoxantha* reference genome using tissue-specific transcriptomic data is therefore essential not only for accurate differential gene expression analyses, but also for reassessing the functional significance of the candidate adaptive variants identified through population genomics (*12*, *14*, *15*, *21*).

Here we assessed gene expression patterns from growing feather follicles across five *Sporophila* species and three plumage patches of different colors. Capuchinos offer the methodological advantage of having low genome-wide differentiation, allowing us to compare species of different colors in a scenario where species differences are otherwise minimal. With these data we tested whether candidate pigmentation genes, which lie in the most highly differentiated regions of the Capuchino genomes, are differentially expressed among growing feather follicles of different colors. Characterizing gene expression across patches, colors, and sexes, we find that these coloration genes are largely not differentially expressed. Gene expression is instead structured primarily by body region. Among coloration genes, only *ASIP* is downregulated in the crown. Moreover, the carotenoid-degrading gene *BCO2* shows a strong male bias that tracks feather carotenoid content. An improved annotation of our reference genome recovered additional genes and previously unknown isoforms and untranslated regions. Under this improved annotation, a subset of pigmentation-associated variants previously classified as noncoding now fall within transcribed sequence, in coding exons and, predominantly, untranslated regions. The absence of expression differences in the candidate outlier genes, together with the recovery of species diagnostic variants within their transcribed regions led us to revise our hypothesis of how coloration differences are shaped in this radiation. We propose that variants in the 5′ and 3′ untranslated regions of transcripts could act post-transcriptionally, contributing to generating coloration differences among species without altering transcript abundance.

## RESULTS

### Transcriptome sequencing and genome annotation

We obtained short-read sequencing data for 43 feather-follicle samples from 29 individuals of five species, including five females. We sampled crown, breast and back plumage patches of five colors (Fig. 1A, Table S1). RNA from nine of these samples spanning all sexes, patches and colors, was pooled for long-read sequencing to improve the *S. hypoxantha* reference genome annotation (Fig. 1B).

**Figure 1.**
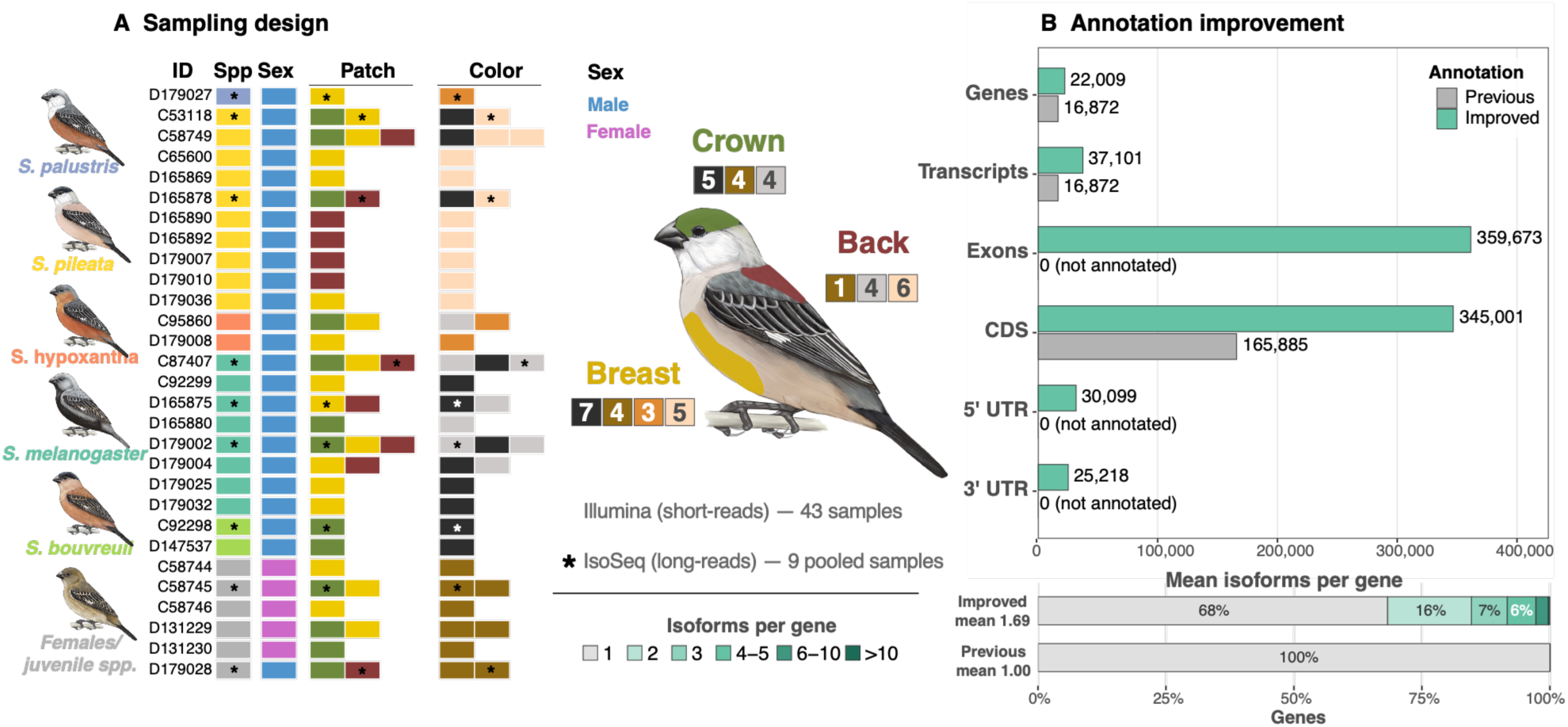
Sampling design and genome-annotation improvement. **(A) Sampling design**. Feather-follicle samples (RNA-seq libraries sequenced with Illumina short-reads; 43 libraries from 29 individuals across five species) are shown with one row per individual. Colored cells indicate individual, species, sex, and the plumage patch(es) sampled (crown, breast, back) together with the corresponding plumage color for each patch. Asterisks (Spp., patch and color columns) mark the nine samples additionally sequenced with PacBio Iso-Seq long-reads. The schematic bird shows the three sampled patches in false color. Colored squares beside each patch show the plumage colors sampled for each patch, with the number of samples for that patch–color combination. **(B) Annotation improvement.** Feature counts for the improved (green) genome annotation, showing gains for different feature types compared to the previous (gray) annotation. The lower panel shows the distribution of the number of isoforms per gene with the percentage of genes in each class color-coded.

This improved annotation increased gene-model resolution relative to the previous one (*15*), recovering 5,137 additional genes (22,009 in total), with the mean number of transcripts per gene rising from 1 to 1.69 (totaling 37,101 transcript models, a 2.2-fold increase). Most genes (15,032; 68.3%) were represented by a single model, whereas 6,977 (31.7%) had two or more isoforms. Coding sequences increased in proportion, from 165,885 to 345,001 CDS features (2.1-fold), now resolved into 359,673 exons and flanked by 5′ and 3′ UTRs (Fig. 1B). Annotation completeness also improved, with BUSCO completeness rising from 95% to 96.9% and missing BUSCOs falling from 4.1% to 2.3%.

### Differential gene expression between feather patches

Global transcriptome structure reflects primarily feather patch, with crown samples separating from breast and back samples along PC1 irrespective of color, while breast and back largely overlapped (Fig. 2A). Additional PCs did not differentiate colors. To identify the genes underlying this structure we fitted an additive generalized linear model (negative binomial, log link) to the counts for each gene (∼ sex + color + patch; hereafter the primary model). We estimated patch effects in a single model that also included sex and color, so that expression differences arising from variation in patch sex and color composition were controlled for, and subsequently tested pairwise contrasts between patches. An ordination restricted to the 121 genes differentially expressed in at least one patch contrast separated crown from breast and back along PC1, with PC2 distinguishing breast and back (Fig. 2B), indicating that the breast–back difference is smaller relative to the overall transcriptome variation (PC1 49% vs. PC2 9%).

**Figure 2.**
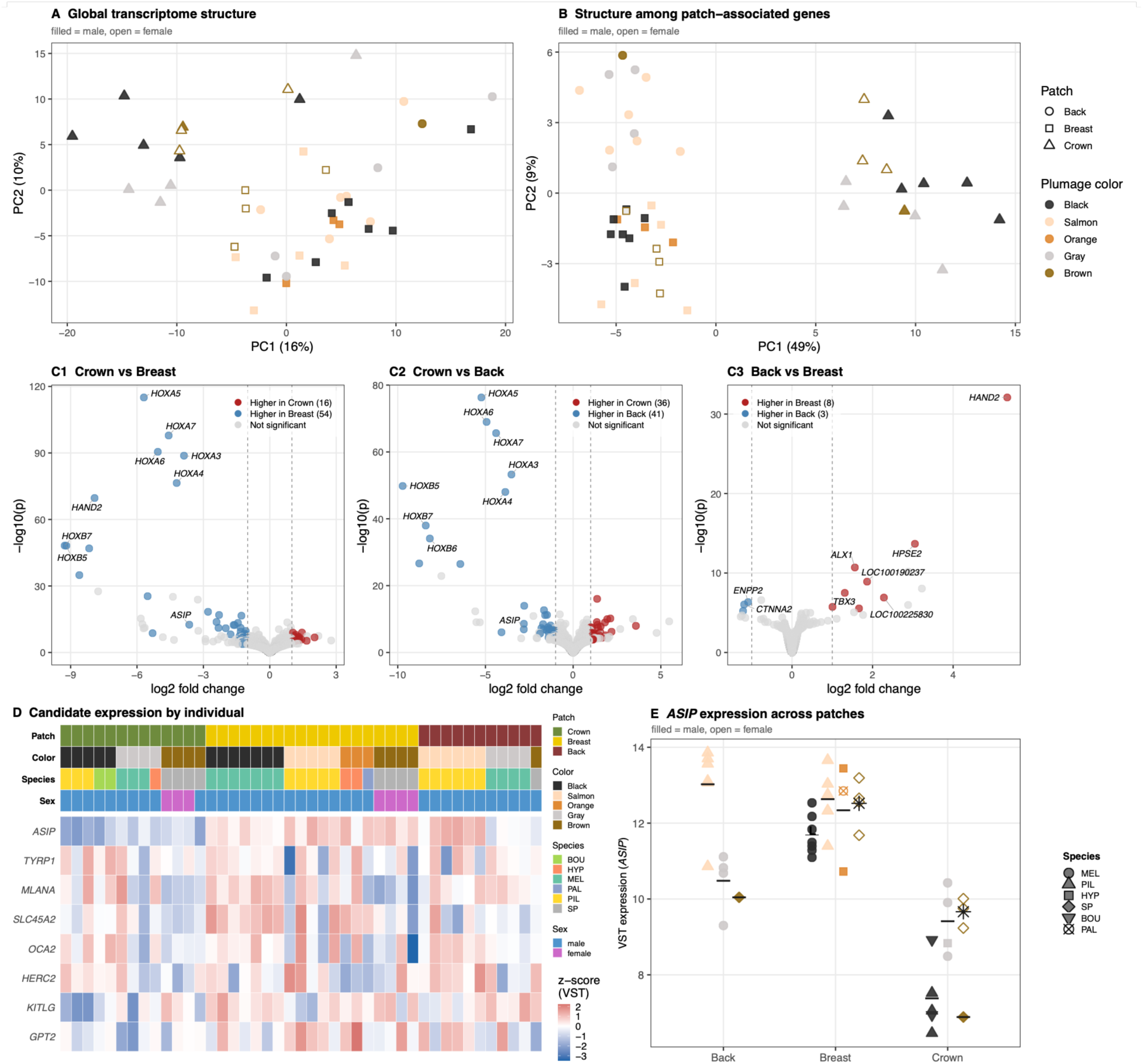
Transcriptomic differentiation among feather patches. **(A)** Principal-component analysis of all 43 samples on the blind (intercept-only) variance-stabilized counts, showing global transcriptome structure. **(B)** PCA restricted to the 121 genes differentially expressed in at least one of the three pairwise patch contrasts extracted from the primary model (∼ sex + color + patch), computed on the same blind variance-stabilized counts. PC1 separates crown from breast and back and PC2 separates breast from back. **(C)** Volcano plots for the three pairwise patch contrasts under the primary model, with Wald tests and ashr-shrunken fold-changes. Crown vs. breast **(C1)**, crown vs. back **(C2)**, back vs. breast **(C3)**. Colored points are differentially expressed genes and the total count per direction is given in parenthesis in each panel. **(D)** Expression of candidate (mostly pigmentation) genes across individual libraries (columns) shown by z-scores per gene on the blind variance-stabilized counts. The annotation strips above the heat map indicate patch, plumage color, species, and sex. Columns are ordered by patch (crown, breast, back), then color and species. **(E)** *ASIP* expression (blind variance-stabilized counts) across the three patches with symbol shape denoting species, fill denoting plumage color, open symbols showing females, and bars marking per-patch male means (asterisk, female mean). Fold-change sign throughout follows the contrast order stated (e.g. crown vs breast: negative = higher in breast). n = 43 libraries from 29 individuals. Species codes: BOU, *S. bouvreuil*; HYP, *S. hypoxantha*; MEL, *S. melanogaster*; PAL, *S. palustris*; PIL, *S. pileata*; SP, brown individuals not assignable to species by plumage.

Differential expression among the three feather patches was pronounced and asymmetric: the crown differed most from the other two patches, with 70 genes differentially expressed between crown and breast and 77 between crown and back, whereas only 11 distinguished breast from back (Fig. 2C, Table S2). Across the three contrasts, 121 genes were differentially expressed in total, with 35 detected in two contrasts and one (*HAND2*) detected in all three (Fig. S1A). Most of these shared genes were associated with the crown contrasts (Fig. S1A), and the main signal came from developmental patterning genes. The strongest gene expression differences included *HAND2* and several *HOX* genes (*HOXA2*–*A7*, *HOXA9*, *HOXB5–B7*, *HOXC6*, *HOXD4*), each expressed at lower levels in the crown and significant in both crown contrasts (Fig. 2C, S1B, Table S3).

Among our candidate pigmentation genes, *ASIP* was the only gene showing differential expression related to patch (as tested by the primary model), with differences between crown and breast (log2 fold-change = −3.64, adjusted *p* = 2 × 10⁻¹⁰) and crown and back (−2.81, adjusted *p* = 3 × 10⁻⁵), but not between breast and back, mirroring the overall crown-driven pattern (Fig. 2D). Crown feather follicles expressed *ASIP* at lower levels than breast or back follicles (Fig. 2E), a difference that held across all sampled species and for both sexes, with female means falling within the male range at each patch (despite their uniformly brown coloration). Breast and back showed statistically indistinguishable higher *ASIP* expression. This patch effect can also be separated from plumage color where the same color was sampled in more than one patch. Black and brown feather follicles, sampled in both the breast and the crown, expressed *ASIP* at higher levels in the breast than in the crown (Fig. 2E). Gray follicles, sampled in the back and crown, showed a weaker difference in the same direction. Salmon-colored follicles, sampled only in the breast and back, showed uniformly high *ASIP* expression, consistent with these two patches being transcriptionally similar. The crown reduction in *ASIP* expression therefore tracks body region rather than plumage color, showing that black plumage color can be generated with both low (crown) and high (breast) *ASIP* expression.

The patch-related transcriptomic signal was robust to model specification (representing different study designs) and to the non-independence of within-individual sampling (Table S3). The 20 most differentially expressed genes were recovered by all models, and 55 of 121 genes (45.5%), were differentially expressed in every model. The remaining genes were those with weaker statistical significance levels and dropped out primarily under models that subset the data and therefore had smaller sample sizes.

### Color-associated gene expression

We sampled plumage patches so that colors could be compared both within and between patches; however, patch and color are confounded in some comparisons. Each patch in our sampling contains a different, unbalanced subset of colors (Fig. 1A), a limitation that we addressed by using several modeling designs. Overall, the transcriptome-wide ordination (Fig. 2A) does not cluster samples by color, and the different model-based tests confirm this result. The number of differentially expressed genes related to color differences was modest under different modeling designs, reaching a maximum of 16 genes in the salmon vs. gray contrast, when controlling for patch differences (∼ patch + color), and 13 when we did not control for co-variates (∼ color; Table S4). None of the *a priori* pigmentation candidates were differentially expressed when patch was included in the model (Table S5). We focused the remaining analyses on the candidate genes in within-patch contrasts.

Restricting comparisons to within a single patch recovered few differentially expressed genes, up to a maximum of 12 (Table S4). From the *a priori* melanogenesis candidates identified as outliers by previous population genomic studies, only two genes were differentially expressed in within-patch color comparisons: *ASIP* and *DCT* (*ASIP* log2 fold-change 2.51, adjusted *p* = 7×10⁻⁶; *DCT* 2.80, adjusted *p* = 1×10⁻³) (Table S5). Otherwise, our candidate coloration genes located in the genomic regions which most clearly differentiate these species, are not differentially expressed in any of our remaining comparisons.

### Sex-biased gene expression and *BCO2*

We found 68 (out of 15,760) genes differentially expressed between sexually dimorphic females and males (Fig. 3A). Most of this signal reflected sex-chromosome dosage: 45 of these genes were Z-linked, all with higher expression in males (Fisher’s exact test, OR = 44.73, *p* = 1.21 × 10⁻⁴⁴; Tables S6 and S7). Across all Z-linked genes we found a median female-to-male expression ratio of 0.67 vs. ∼1.00 for autosomal genes (Wilcoxon *p* = 1.54 × 10^⁻197^; Fig. S2A), indicating partial dosage compensation in heterogametic females. From the 23 autosomal genes with sex-biased expression patterns, ten had higher expression in males and thirteen in females (Fig. S2B).

**Figure 3.**
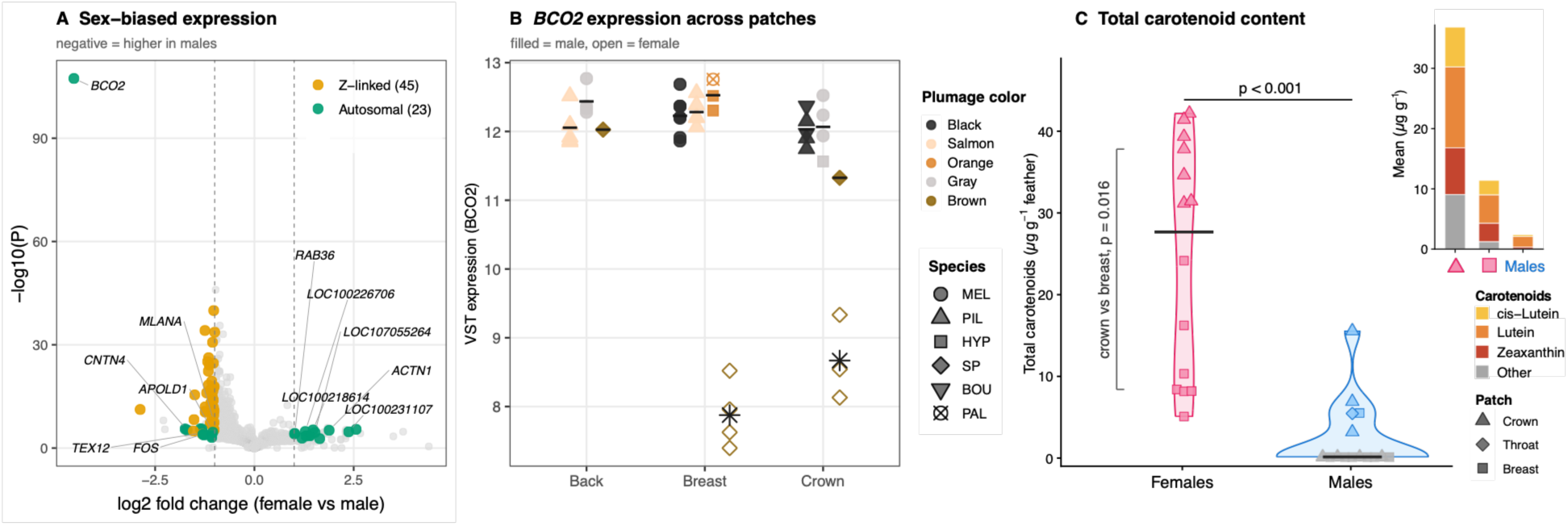
Sex-biased expression, *BCO2*, and feather carotenoid content. **(A)** Volcano plot of female-versus-male differential expression across the 15,760 tested genes (primary model ∼ sex. Wald test with apeglm-shrunken fold-changes for the two-level contrasts, see Methods). Significant genes are colored by chromosome-linkage class (Z-linked, autosomal). The ten autosomal genes with the strongest sex-bias, *FOS*, and *MLANA* are labeled. **(B)** *BCO2* expression (blind variance-stabilized counts) across the three feather patches. Details as in Figure 2. **(C)** Total feather carotenoid content (µg g⁻¹) in females and males. Violins show the distribution, points are individual samples shaped by feather patch, and the crossbar indicates the median. Gray points mark male samples below the detection threshold of 2 µg g⁻¹. The horizontal bracket shows a two-sided Wilcoxon rank-sum test between sexes, while the vertical bracket gives a paired Wilcoxon test of crown versus breast within females (n = 7, each sampled at both patches). The sex difference was consistent across species, though sample sizes were too small for species-level comparisons. Inset: mean concentration of each carotenoid in female crown, female breast, and male feathers. Carotenoid samples comprise 29 feather samples from 15 individuals of seven species, including three species (*S. hypochroma*, *S. iberaensis* and *S. ruficollis*) not represented in the RNA-seq dataset. n = 43 libraries from 29 individuals in (A–B).

The largest effect of any gene belonged to the carotenoid-cleavage gene *BCO2* (log2FC = −4.55, adjusted *p* = 5.95 × 10⁻¹⁰⁴; ∼23-fold higher in males; Fig. 3A, Table S7), which had a 1.8-fold larger effect than that of the next most strongly differentially expressed autosomal gene. *BCO2* was significant in all models, including those controlling for patch and color (Table S7), and in models that subsampled the dataset. The higher *BCO2* expression in males was consistent across the three feather patches (Fig. 3B). The single brown-plumaged juvenile male (D179028) resembled other males rather than brown females for *BCO2* expression (Fig. 3B). Consistent with *BCO2*’s role in carotenoid degradation, total feather carotenoid content was markedly lower in males vs. females (Fig. 3C, Table S8).

Because several individuals contributed samples from more than one patch, we repeated the sex contrast on one randomly chosen sample per individual, choosing 20 different times. This separated robust from weakly supported genes. When comparing females and males without accounting for body region (∼ sex, the uncontrolled design), and again with body region included as a covariate (∼ patch + sex, the patch-adjusted design), two of the 23 autosomal genes were significant in every patch-adjusted draw (*BCO2* and *FOS*), seven in at least half, and five in none. *FOS* reached significance in all 20 patch-adjusted draws but in none of the unadjusted draws, so its sex effect emerged only once variation among patches was accounted for. Differences in the expression of Z-linked genes were largely robust (35 of 45 significant in at least half of the patch-adjusted draws), consistent with a chromosome-wide dosage effect.

Among candidate pigmentation genes, only the expression of *MLANA* was sex-biased, but it is Z-linked with a female-to-male expression ratio of 0.47, consistent with an absence of dosage compensation. The other two Z-linked candidates, *SLC45A2* and *TYRP1*, had ratios of 0.62 and 0.79, close to the median for all Z-linked genes (0.67).

### An improved genome annotation reclassifies candidate outlier variants into coding (CDS) and untranslated regions (UTR)

Our improved genome annotation shows that a fraction of our outlier markers (SNPs and indels) are located within transcribed sequence, primarily within untranslated regions. Of 588 unique outlier positions associated with candidate melanogenesis genes, which lie in the most strongly differentiated regions of the genome (see Methods), 56 gained a CDS or UTR annotation absent from the previous genome annotation: 5 were reassigned to CDS and 18 and 33 to 5′ and 3′ UTR, respectively (Fig. 4A). The same pattern held genome-wide: of the 10,762 outlier positions across the genome, 470 moved from noncoding into primarily UTR and CDS, while also 28 previously coding positions were reassigned to 5′ UTR (6) or noncoding (22) sequence (bringing the totals in these classes to 378 CDS, 223 5’ UTR and 148 3’ UTR; Fig. S3). SNPs and small indels were reclassified at similar rates (4.4% and 4.5%). The 41 outlier SVs (12 deletions, 27 insertions, 2 complex, with a median length of 155 bp, and a maximum length of 6.3 kb) all remained intergenic or intronic (median distance to the nearest CDS 15.6 kb).

**Figure 4.**
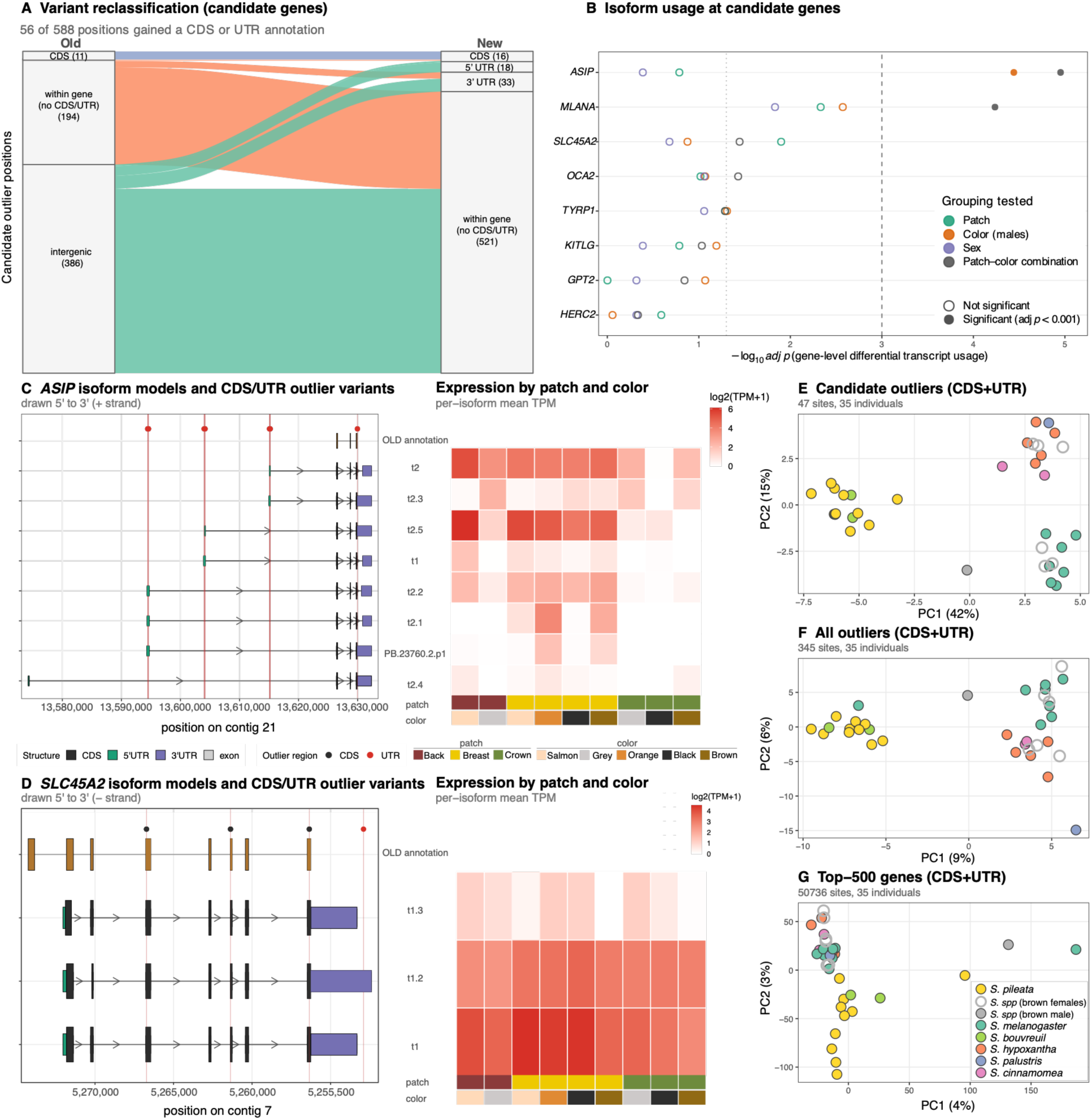
Reannotation of pigmentation-associated outlier variants, isoform usage, and transcript-derived genotype clustering. (**A**) Reclassification of candidate outlier positions under the previous and improved genome annotations. (**B**) Differential transcript usage (DTU) at candidate pigmentation genes, testing whether the relative proportions of a gene’s isoforms differ among groups independently of the gene’s total expression. Each row is a candidate gene with at least two expressed isoforms, and each point is one DTU test, colored by the grouping tested: feather patch (crown, breast and back; 43 samples), plumage color in males (34 samples; females were excluded because all sampled females are brown), sex (43 samples), and the combination of patch and color (42 samples). The dotted line marks adjusted *p* = 0.05 and the dashed line adjusted *p* = 0.001 with filled circles showing tests passing our statistical significance threshold. (**C**, **D**) Gene models for *ASIP* and *SLC45A2* showing exons, CDS and UTRs for expressed isoforms in the improved annotation, alongside the single model from the previous annotation, with vertical lines marking outlier variants overlapping CDS or UTR regions. Heatmaps give mean per-isoform abundance (log_2_ TPM + 1) across plumage patch–color conditions. A fourth *SLC45A2* model, expressed at low levels (2.9% of gene expression) and with a reading frame shifted relative to the other isoforms, was included in the differential transcript usage tests but is not drawn. (**E** to **G**) Principal component analyses of 35 individuals based on CDS and UTR genotypes at candidate outliers (47 sites), all outliers (345 sites), and variants in the 500 most highly expressed genes after removing those overlapping candidate genes (50,736 variable sites). Genotypes were called from RNA-seq reads. Points are colored by species with open circles denoting brown individuals that cannot be assigned to species by plumage.

Functional consequence prediction showed that the candidate outlier variants falling within annotated CDS included both synonymous and missense changes. Of the 588 outlier positions within candidate genes, 16 fall in coding sequence: eight are missense changes, seven synonymous and one splice-site variant within a coding exon. Two further splice-site variants lie just outside coding exons (Table S9). The missense substitutions occur mostly in genes encoding melanosomal proteins, one each in *TYRP1* and *MLANA*, and two each in *OCA2* and *SLC45A2* (additionally, one each in *GPT2* and *HERC2* which are not involved in melanin regulation). Allele frequencies at these positions, obtained from the whole-genome data underlying the F_ST_ and GWAS outlier scans (*15*), showed that none of these changes are private to a single species. The *SLC45A2* substitution (Asn368Asp) reaches high frequency in *S. hypochroma* (0.86), *S. melanogaster* (0.81) and *S. ruficollis* (0.80), is absent from *S. bouvreuil*, *S. nigrorufa* and *S. palustris*, and segregates at low to intermediate frequency (0.07 to 0.43) in the remaining four species. The *TYRP1* substitution (Ser443Pro) is most frequent in *S. ruficollis* (0.83) and *S. iberaensis* (0.79), absent from *S. melanogaster*, and segregating at low to intermediate frequency (0.05 to 0.34) in other species. This substitution, however, is predicted only on a minor isoform accounting for 0.4% of *TYRP1* expression in feather follicles. The two *OCA2* substitutions, which the previous gene model had not placed on the protein (*14*), resolve to Arg8Lys and Met126Thr and differ among species in opposite directions: Arg8Lys is frequent in *S. palustris* (0.87), *S. nigrorufa* (0.83) and *S. hypoxantha* (0.69) and rare in *S. iberaensis*, *S. melanogaster* and *S. pileata*, whereas Met126Thr shows the reverse pattern. The *SLC45A2* change at Asn368Asp corresponds to the exonic substitution reported by Estalles et al. (*14*), which the improved gene models place at the same residue. Because these positions were ascertained as outliers, differentiation among species is expected; the relevant observation is that the alternate alleles are shared across species in different combinations rather than confined to one. Beyond SNPs, a 1 bp deletion was reclassified from intergenic to the *KITLG* 5′ UTR, and 1 and 5 bp insertions were reclassified from intergenic to a *GPT2* 3’ UTR. Notably, for *ASIP*, the candidate with the strongest differential expression signal, all 230 of its outlier variants fell in its 5′ UTRs (14), 3′ UTR (1), or introns (215). Eleven of the fourteen 5′ UTR variants are fixed for opposite alleles between groups of species, and at two additional variants, alternate allele frequencies differ by at least 0.95 between species (Table S9).

### Differential transcript usage is limited at candidate coloration genes

Given that the differential expression of candidate coloration genes does not explain color differences, we asked whether a shift in isoform use could modulate pigment production and shape feather coloration. The improved annotation resolved multiple isoforms at candidate loci that the previous annotation lacked, allowing relative isoform usage to be tested directly. Across the eight candidate genes with at least two expressed isoforms, only three of 32 tests showed differential transcript usage (adjusted *p* < 0.001), and all involved color-related comparisons. *ASIP* isoform usage differed across colors in males (adjusted *p* = 4 × 10⁻⁵) and both *MLANA* (adjusted *p* = 6 × 10⁻⁵) and *ASIP* (adjusted *p* = 1 × 10⁻⁵) showed differences in one comparison between patch–color combinations (Fig. 4B, Table S10). We did not find differential isoform usage between sexes or plumage patches. For *MLANA* the switch in the dominant isoform usage was small (1.1 percentage points between patch–color groups), despite being statistically significant. *ASIP* was the only gene showing a larger change in isoform composition (Fig. S4).

The isoform structure underlying these tests is illustrated by *ASIP* and *SLC45A2*, for which we identified eight and four expressed isoforms differing in exon and untranslated-region structure, respectively. In *ASIP*, 15 of the outlier variants fall within these newly resolved transcript features, whereas all but one of the *SLC45A2* outlier variants overlap coding sequence (Fig. 4C, 4D). In *SLC45A2*, isoform t1 predominated in every patch–color condition (48 to 67% of the gene’s expression), followed by t1.2 (23 to 43%), whereas t1.3 contributed 6 to 13%. In *ASIP*, by contrast, dominance alternated between two isoforms, t2.5 and t2, depending on the patch–color combination (Fig. 4C, 4D, right).

Differential isoform usage is therefore not a general mechanism across all our candidate genes for shaping color variation, but shows substantial variation in *ASIP*. Across patch–color combinations, the *ASIP* isoform t2.5 accounts for 62% of the gene’s expression in salmon back feathers and 60% in salmon breast feathers, but only 10% in gray back feathers, where t2 takes over as the dominant isoform. Gray crown feathers show the same pattern, although *ASIP* is expressed at low levels in the crown and these estimated proportions are based on few reads (Fig. 4C, Fig. S4). Within the breast, where all four colors are represented and coverage is highest, the variation largely disappears: t2.5 accounts for 42 to 60% of *ASIP* expression in black, brown, orange and salmon feathers alike. Therefore, gray feathers use the same *ASIP* isoform composition in both regions where they occur, yet within the breast the four remaining colors converge on similar proportions (Fig. S4). Isoform usage at *ASIP* could separate gray from the other colors rather than distinguishing colors generally, yet better sampling is needed to resolve this difference robustly.

### Candidate-gene outlier variants in coding and untranslated regions of transcripts can be used to identify species

Given the pattern of patch-specific but not color-specific gene transcription, we asked what information is carried on the transcripts themselves, where variants could act post-transcriptionally rather than through generating differences in transcript abundance. We therefore genotyped outlier variants identified from whole-genome data directly from RNA-seq reads and tested whether they can be used to differentiate species. Individuals clustered by species in a principal component analysis of 35 individuals genotyped at candidate-gene CDS and UTR outlier variants: the 29 individuals used for the expression analyses and six additional birds (11 samples) from a pilot study not included in the differential expression analysis (Fig. 4E-G, Fig. S5).

In the combined analysis with CDS and UTR variants, individuals clustered according to species, and females and an immature brown male, which cannot be assigned to species by plumage, fell within those same clusters, suggesting these individuals belong to the same species as the adult males (Fig. 4E-G, Fig. S5). Clustering and separation were more pronounced than for the full set of F_ST_/GWAS outliers, which includes singleton and weaker statistical associations from SNPs falling outside the main divergence peaks, and weaker still for all variants from the 500 most highly expressed genes excluding candidate genes. The strength of the clustering was mirrored by the sum of the variance explained by PC1 and PC2: 57% for candidate-gene outliers, 15% for the full outlier set, and 7% for the 500 most highly expressed genes (Fig. 4E-G). This stronger separation was recovered despite the candidate set containing roughly 7-fold and 1,000-fold fewer variants than the broader outlier and top-500 sets, respectively.

The same pattern held when candidate-gene outliers were partitioned by sequence class. Both the 15 CDS and the 32 UTR variants alone could be used to cluster individuals by species (Fig. S5), implying that coding and untranslated variants within candidate pigmentation genes carry a species-specific signal. Species-diagnostic information is present on the mature mRNA, where it could act to shape coloration downstream.

## DISCUSSION

The genomic regions distinguishing Capuchino Seedeater species are few and narrow, and most contain or are adjacent to pigmentation genes. Because most differentiated variants in these regions were thought to be noncoding, male species-specific plumage patterns were hypothesized to arise from changes in the expression of these otherwise mostly conserved candidate genes (*12*, *14*, *15*). Sampling growing feather follicles across species, plumage patches, and colors, we instead found that feather follicle transcriptomes were structured by plumage patch and by sex, and that candidate pigmentation genes were predominantly not differentially expressed among colors. We discuss these results in turn and propose a model in which species-diagnostic variants carried on pigmentation-gene transcripts act within the context of each plumage patch to produce species-specific plumage patterning.

### Plumage patches have distinct transcriptomic signatures independent of color

The most robust differential gene expression signal in our data was positional. Crown follicles were transcriptionally distinct from breast and back follicles, which in turn shared a largely similar profile, a pattern driven primarily by the downregulation of *HAND2* and several *HOX* genes in the crown. *HOX* genes provide the positional information in the dermis that shapes the development of epidermal appendages (*22*, *23*), and a similar *HOX* and *HAND2* signature distinguishes body regions in Dark-eyed Juncos (*Junco hyemalis*) (*24*). Additionally, *ZIC1*, a gene regulating feather formation in chickens (*25, 26*), was upregulated in back follicles. *ALX1*, a gene with expression patterns that mark color boundaries in Estrildid finches (*27*), showed higher expression in Capuchino crown and breast follicles. Both genes also show a dorsoventral expression pattern in domestic Duck skin (*28*). Follicles in different plumage patches therefore carry distinct transcriptomic signatures, independently of their pigmentation. This is further illustrated by the expression of the melanogenesis regulator *ASIP*, which was downregulated in crown follicles in every species sampled and in both sexes. Moreover, follicles of the same color differed in *ASIP* expression depending on the patch in which they grew: black and brown follicles expressed less *ASIP* in the crown than in the breast. Spatially restricted *ASIP* expression is a recurrent mechanism of vertebrate color patterning (*10*). In birds, differences in *ASIP* expression among plumage regions have been described in chickens, where expression from the distal promoter occurs in ventral plumage and in gold ornamental feathers on the back of males, as well as in the ventral plumage of Black Sparrowhawks (*Astur melanoleucus*) (*29*, *30*).

Two other genes with patch-associated expression patterns are known to mediate pigment deposition. *TBX3*, enriched in breast follicles, is not a canonical pigmentation gene, yet cis-regulatory mutations affecting its follicular expression underlie the Dun camouflage pattern of horses, through a reduction in *KITLG*, restricting where pigment is deposited within the hair (*31*). *CORIN*, which encodes a protease that attenuates Agouti signaling in mouse hair follicles (*32*, *33*), was reduced in the crown alongside *ASIP*, and both genes also differ among body regions in uniformly black Carrion Crows (*Corvus corone*) (*34*). Each patch therefore offers a different transcriptomic context, including genes that shape how pigment is deposited, in which the candidate pigmentation genes identified as divergent in Capuchinos are expressed.

In Capuchinos, crown melanosomes are larger and more elongated than those of other patches, and crown feathers are almost entirely eumelanic, producing both black and gray crowns (*35*). Reduced *ASIP* expression weakens antagonism of *MC1R* and favors eumelanin synthesis, and eumelanosomes contain a fibrillar *PMEL* scaffold that permits elongated shapes, whereas pheomelanosomes remain spherical (*36*). Low crown *ASIP* expression could therefore determine the eumelanic crown common to all males in the radiation. The single exception to crown melanosome morphology is in *S. iberaensis*, which has round crown melanosomes despite their high eumelanin content (*35*). Our current dataset does not include samples from this species’ crown, which would allow us to assess if *ASIP* expression differs from that of the remaining species with elongated crown melanosomes. Overall, these morphological and transcriptomic results suggest that the same colors may not be produced by the same mechanisms in different plumage patches.

### Candidate pigmentation genes are not differentially expressed among plumage colors

Capuchino species show very low genome-wide differentiation (F_ST_ ∼0.008) (*12*), and against this homogeneous background candidate pigmentation genes lie in the few narrow regions that consistently differentiate species. In some cases, these are the only regions of the genome that allow species to be diagnosed genetically (*12*, *14*, *15*). Under the previous genome annotation, approximately 97.2% of outlier variants fell outside annotated coding sequence (*15*), and many did so within conserved genomic regions across several bird species (*12*), which suggested that they could fall within cis-regulatory elements (e.g., enhancers) that alter the expression of the nearby pigmentation genes (*12*, *14*, *15*). Our data do not support this expectation in growing feather follicles as most candidate gene expression did not differ among colors (except for *ASIP* and *DCT* expression between salmon and gray plumage, when restricted to back follicles). Isoform usage at candidate genes was similarly stable with statistically significant differences only changing the proportion of the dominant isoform substantially for *ASIP*. However, that shift is not a general color effect. It is likely driven by gray feathers, in which the expression of the dominant isoform is replaced by a different transcript, a pattern repeated in both the crown and the back. Within the breast, where four colors (but not gray) occur and coverage is highest, isoform proportions are similar. Overall, the expression of these pigmentation genes or their isoform usage does not track plumage color.

This lack of differential gene expression at pigmentation genes contrasts with other avian systems with melanic plumage, which similarly sampled growing feather follicles from captive and wild birds. In Dark-eyed Juncos, plumage color differences are associated with the expression of *ASIP* and other pigmentation genes in the absence of coding variation (*24*), and in Black Sparrowhawks, plumage morphs are strongly associated with *ASIP* expression again without morph-associated coding variation in *ASIP* or *MC1R* (*29*). Capuchinos show a different pattern where candidate loci are differentiated in primarily noncoding sequence but without color-associated transcript abundance. Our findings show that variants acting to change pigmentation-gene expression are not the main route to plumage divergence in this system. Intergenic outliers nonetheless could still modify gene expression in plumage patches or species that our sampling of growing follicles did not capture (e.g., the throat patch in *S. ruficollis* or *S. palustris*).

This absence of color-associated differential gene expression is unlikely to reflect sampling the wrong tissue or cell type, or the loss of a color-specific transcriptomic signal from experimental noise. Melanocytes determine both the type and the amount of pigment deposited, and they are a minority population within the feather follicle, so a bulk transcriptome could in principle dilute their signal. In our libraries, however, canonical melanocyte markers are among the most highly expressed genes in the dataset: *TYRP1*, *PMEL* and *MLANA* rank in the top 1% of all genes by mean normalized expression, and *EDNRA*, *MITF*, *KIT* and *TYR* in the top 15%, with expression of these genes detected in all of our samples. *MLANA* and *PMEL* are melanocyte-specific markers in single-cell data from developing chicken feathers (*37*). The follicles we sampled during active feather growth therefore contain the pigment-producing cells. The same samples also recover a large and consistent difference in *ASIP* expression between body regions, confirming that our study design can detect expression differences at pigmentation genes.

### Sex-biased *BCO2* expression depletes dietary carotenoids in male plumage

Previous work in this system has focused on the melanin-based plumage of adult males, a trait that differs among species and mediates assortative mating (*12*, *14*, *21*). Sampling both sexes revealed an additional, sex-specific pigmentary difference. *BCO2*, which encodes the carotenoid-cleaving enzyme β-carotene oxygenase 2 (*38*), was the gene with the strongest sex-biased expression (∼23-fold higher in males). Consistent with this result, feather carotenoid analyses of seven females from five species, three of which were not represented in our expression dataset, revealed that lutein and zeaxanthin were the predominant carotenoids and occurred at higher concentrations in the crown than in the breast. In contrast, carotenoids were undetectable in most male samples (10 of 14) and present only at low concentrations in the rest. Because birds cannot synthesize carotenoids, and deposit these yellow dietary xanthophylls in feathers after acquiring them from their diet without metabolic modification (*39*), the sex difference likely reflects carotenoid degradation within the follicle, the step at which *BCO2* acts.

This inverse relationship between follicular *BCO2* expression and feather carotenoid content parallels what has been found in other birds. In canaries and Northern Cardinals (*Cardinalis cardinalis*), higher *BCO2* expression in developing female feathers degrades carotenoids and produces paler plumage (*40*, *41*). In Red-backed Fairywrens (*Malurus melanocephalus*), *BCO2* is upregulated in the carotenoid-free cream feathers of unornamented males compared to the black feathers of ornamented males (*42*). In these examples high *BCO2* expression is seen in the less ornamented sex/individuals, whereas in Capuchinos carotenoid degradation occurs in the ornamented sex. The single brown-plumaged male in our dataset expressed *BCO2* at levels similar to other males, as expected if it was molting into definitive plumage and its growing follicles were removing carotenoids.

Carotenoid removal in males and the retention of these pigments in females may serve different functions. In males, degrading dietary carotenoids could keep the melanin-based colors of the breeding plumage free of yellow tones that would alter the species-specific melanin-based signal. In females, retained lutein and zeaxanthin may function in crypsis. Together with studies of species with carotenoid-based male ornamented feathers, where high *BCO2* expression produces drabber females (*40*, *41*), our results suggest that sexual dichromatism can arise from either up-or downregulation of *BCO2*.

### Species-diagnostic information is carried on pigmentation-gene transcripts

If candidate genes are not differentially expressed among colors, their differentiated sequence could influence pigmentation in other ways. Our improved annotation shows that part of this sequence lies on the transcripts themselves. Genome-wide, adding UTRs and additional transcript models raised the proportion of outlier variants within coding or untranslated sequence from 2.8% to 7.0%, and within the candidate genes 51 of the 56 positions newly assigned to transcribed sequence fell in UTRs. Genotypes at coding and UTR variants in candidate genes, called directly from follicle RNA-seq reads, clustered individuals by species. This reclassification places species-diagnostic information on the mature mRNA of pigmentation genes, where it could affect pigment production without changing transcript abundance.

UTR variants, which in Capuchinos account for most of the differentiated variants on the mRNA, can have regulatory properties and measurable phenotypic effects have been described in humans (*43*). Such variants may alter translation, mRNA stability, localization, and processing, through upstream open reading frames and RNA structure in 5′ UTRs and through microRNA and AU-rich binding sites and alternative polyadenylation in 3′ UTRs (*44*, *45*). UTR variants can also create or disrupt sites of N⁶-methyladenosine (m6A), the most abundant mRNA modification, which is enriched in 3′ UTRs and near stop codons and regulates mRNA decay and translation (*46*, *47*). Effects on translation or localization alter protein output without changing mRNA abundance and are therefore undetectable through RNA-seq (*48*, *49*). All outlier variants at *ASIP*, a dose-sensitive antagonist of melanocortin signaling (*10*), fall in UTRs or introns. *ASIP* is also the one candidate with expression levels that vary with plumage color, differing between salmon and gray backs. UTR variants could change transcript stability and therefore abundance, or change translation and protein output alone, so both routes are possible at this locus. Missense substitutions also occurred in genes encoding melanosomal (*TYRP1*, *OCA2*, *SLC45A2*, and *MLANA*) and other (*HERC2* and *GPT2*) proteins, and some were largely restricted to particular species, such as the *TYRP1* substitution frequent in *S. ruficollis* and *S. iberaensis*, but absent from *S. melanogaster*. Such changes could modify protein activity, stability, or localization. Within-individual variation, however, cannot be explained by coding differences as each bird carries the same variants in every follicle. What differs between patches is the positional context in which those transcripts function.

### A model for patch-specific plumage patterning

Our results suggest that species-specific plumage patterns arise from the interaction between variants carried on candidate pigmentation-gene transcripts and the transcriptional context of each plumage patch. Follicles in different patches have distinct transcriptional identities (e.g., through *HOX*, *HAND2* and *ASIP* expression), whereas candidate pigmentation genes are expressed at similar levels regardless of color. While a species-specific UTR or coding variant is present in every follicle, its effect could depend on the patch in which it is expressed, for example through the RNA-binding proteins and microRNAs available to act on a UTR, or on the level of *MC1R* antagonism under which a melanosomal protein functions. The same allele could therefore contribute to generating different colors in different patches. Variants within these genes lead to different alleles, which could generate different colors in the same patch of different species (e.g., a salmon vs. black breast in two species carrying different alleles at a coloration gene). Our *ASIP* data show that color is not a direct indicator of its expression level, since black follicles occurred both in the crown, where *ASIP* expression was low, and in the breast, where it was high. Intergenic outliers could add cis-regulatory elements active only in particular patches (not sampled here, such as the throat), stages, or melanocyte populations, which would extend rather than contradict this model. The model concerns the patterning of male plumage and is separate from the sex difference in carotenoid content, which involves a different pigment class and likely a different regulatory pathway (involved in coloration differences between males and females).

This model makes testable predictions. Reporter assays and ribosome profiling could test whether candidate-gene alleles differ in translation, stability, or processing, and patch-resolved chromatin accessibility and single-cell transcriptomics across stages of feather growth could reveal cis-regulatory effects that bulk RNA-seq dilutes. Our sampling covered a subset of Capuchino species and three plumage patches at a single stage of feather growth, so differential expression of pigmentation genes may still contribute to color differences in species, patches, or stages we did not examine.

More broadly, our results caution against equating noncoding outlier variants with possible changes in gene expression. In Capuchino Seedeaters, the few genomic regions that distinguish species appear to shape plumage not by changing how much pigmentation genes are expressed, but through sequence differences on their transcripts with effects that depend on the transcriptional context of each plumage patch.

## MATERIALS AND METHODS

### Study system and sampling

We sampled individuals at two migratory stopover sites in southeastern Brazil, Pouso Alegre (Minas Gerais) [22°07’20,3”S; 45°51’27,4”W] and Indaiatuba (São Paulo) [23°09’20,4”S 47°14’18”W] in October/November of 2024, where birds were molting into reproductive plumage. Individuals belonged to six *Sporophila* species: *S. hypoxantha* (Tawny-bellied Seedeater), *S. cinnamomea* (Chestnut Seedeater), *S. melanogaster* (Black-bellied Seedeater), *S. palustris* (Marsh Seedeater), *S. bouvreuil* (Capped Seedeater), and *S. pileata* (Pearly-bellied Seedeater) (Fig. 1A, Table S1). We retained unidentified individuals with brown plumage (females or juveniles of undetermined species, hereafter ‘SP’) as a separate category. We only plucked growing feathers at a comparable early developmental stage, while still completely enclosed in the sheath or just emerging from it (∼1 mm), to sample the distal portion of the feather which varies in color across patches and species. For each individual, we sampled up to three plumage patches: crown, breast, and back. We grouped belly samples with breast given their anatomical proximity and that both patches show the same color in the different Capuchino species. We recorded plumage color for each plumage patch, sampling black, brown, salmon, orange, and gray growing feathers (Fig. 1A).

We also included 11 samples from six individuals collected in a pilot study in Corrientes Province, Argentina: three *S. hypoxantha* (one brown female), two *S. cinnamomea* and one unidentified brown individual (SP) (Table S1). Because RNA from these samples was more degraded, they were processed with a different library preparation protocol (see below) and used only for genotyping, not for expression analyses.

### Illumina RNA sequencing and quality control

Total RNA was extracted from growing feather follicles preserved in RNA/DNA Shield (Zymo Research), pooling on average five feathers per sample (range 2–10), using the Zymo Research Quick-RNA Microprep Kit according to the manufacturer’s protocol. Samples were first homogenized in 2 mL bead mill tubes containing ceramic beads (VWR #10158-612) using a VWR 4-Place Mini Bead Mill Homogenizer (VWR #10158-558), and subsequently processed with the RNA kit, including a Proteinase K treatment and lysis. RNA was purified using silica spin columns, with DNase I digestion performed on-column to eliminate genomic DNA contamination, and eluted in nuclease-free water. We measured RNA concentration with a Quickdrop spectrophotometer (Molecular Devices, San Jose, CA, USA) and assessed integrity with a 5200 Fragment Analyzer using the HS RNA Kit (15 nt; DNF-472, Agilent Technologies, Santa Clara, CA, USA). RNA Quality Numbers (RQN), calculated in ProSize v5.0.1.6 (Agilent), ranged from 7.6 to 10 in the samples used for gene expression analyses (primary libraries) and ranged from 1.1 to 7.1 in the pilot samples.

Primary libraries were enriched for mRNA by polyA selection, whereas pilot libraries were depleted of rRNA with the NEBNext rRNA Depletion Kit. All libraries were prepared with the NEBNext Ultra II Directional RNA Library Prep Kit and sequenced as 2×150 bp paired-end reads on an Illumina NovaSeq X at a minimum target depth of 20 million read pairs per sample. Sequencing was performed by the Transcriptional Regulation and Expression (TREx) Facility at the Cornell University Biotechnology Resource Center (RRID:SCR_022532). We sequenced 46 libraries from 31 individuals in the primary dataset and 11 libraries from six individuals in the pilot dataset.

Raw read quality was assessed using FastQC v0.12.1 and MultiQC v1.31 (*50*, *51*). Reads were trimmed with fastp v0.23.4 (*52*): adapter sequences were detected automatically for paired-end reads (“detect_adapter_for_pe”), and reads shorter than 30 bp after trimming were discarded (“length_required 30”). Trimmed reads were aligned to the *S. hypoxantha* reference genome (HYPOXB009684, GCA_053841625.1) with STAR v2.7.11b (*53*), using an index built with the annotation to incorporate annotated splice junctions and a splice-junction overhang of 149 bp. We used STAR’s two-pass mode (--twopassMode Basic), in which splice junctions discovered in a first pass are added before the final alignment, discarded alignments with non-canonical splice junctions (--outFilterIntronMotifs RemoveNoncanonical), and added the strand attribute (--outSAMstrandField) required by downstream tools to spliced reads, producing coordinate-sorted BAM files.

### Pacific Biosciences Kinnex full-length RNA sequencing

Nine samples encompassing all patches and colors were pooled, and Kinnex full-length RNA libraries were prepared and sequenced on a PacBio Revio instrument at the Weill Cornell Medicine Genomics Resources Core Facility (RRID:SCR_025634). Sequencing generated 10.9 million HiFi reads totaling 168.9 Gb, with a mean read length of 15,496 bp. Median read quality was Q36, with an average of nine passes per molecule and resulting in 8.50 million reads (78.0%) with a quality score ≥Q30. We processed raw HiFi reads through the standard Iso-Seq pipeline (*54*): including segmentation with skera v1.4.0, primer removal and demultiplexing with lima v2.13.0 (--isoseq --peek-guess), polyA tail trimming and chimeric read removal with isoseq refine (--require-polya), transcript clustering and consensus generation with isoseq cluster2, genome alignment with pbmm2 v1.7.0 (--preset ISOSEQ --sort), and transcript collapse with isoseq collapse (isoseq v4.0.0). This produced 769,961 full-length transcript models. Transcript abundance was estimated from full-length (FL) read counts per isoform.

We then filtered these 769,961 transcripts along two independent paths serving different purposes: a stringent filter for coding sequence (CDS) prediction (76,135 transcripts remained; see *Iso-Seq transcript classification and CDS prediction*), and a simple FL-read-depth filter for UTR prediction (104,871 transcripts supported by ≥5 FL reads; see *UTR annotation with PASA*).

### Reference genome annotation

#### Evidence based gene prediction

We used the *S. hypoxantha* reference genome (HYPOXB009684, GCA_053841625.1) as the annotation target with repetitive elements soft-masked, and performed gene prediction with BRAKER3 (*55*) in ETP (Evidence-based Training and Prediction) mode. We supplied all 46 short-read RNA-seq BAM files alongside a combined protein database of OrthoDB v11 Vertebrata sequences and *Taeniopygia guttata* proteins (9,822,322 protein sequences). BRAKER3 internally used GeneMark-ETP for model training and AUGUSTUS for gene prediction, with ProtHint for protein hint generation, producing 19,506 gene models.

#### Iso-Seq transcript classification and CDS prediction

We quality-filtered and structurally classified full-length Iso-Seq transcripts with SQANTI3 v5.2.1 (*56*) relative to the existing *S. hypoxantha* annotation. Of 769,961 input transcripts, 76,135 passed quality filtering, which required canonical splice sites, no reverse transcriptase switching artifacts, and class-specific minimum full-length (FL) read support. We required ≥2 FL reads for full splice match (FSM), incomplete splice match (ISM), and novel in catalog (NIC) transcripts; ≥5 for novel not in catalog (NNC) transcripts; and ≥10 for intergenic transcripts. We predicted coding sequences (CDSs) with TransDecoder v5.5.0 (*57*), retaining the single best ORF per transcript (72,888 ORFs, 50,133 of them complete), and converted them to genomic coordinates with the TransDecoder utility cdna_alignment_orf_to_genome_orf.pl, which propagated 70,722 ORFs across 9,754 loci.

#### Integration and gap-filling

We integrated BRAKER3 gene models and SQANTI3-filtered TransDecoder predictions with TSEBRA (*58*), in its version for long-read data, yielding 20,743 protein-coding gene models. To recover genes captured by neither BRAKER3 nor the Iso-Seq data, we generated homology-based predictions with GeMoMa v1.9 (*59*) using reference annotations from the Zebra Finch (*Taeniopygia guttata*, GCF_048771995.1, bTaeGut7.mat; 23,294 annotated genes) and the chicken (*Gallus gallus*, GCF_016700215.2, bGalGal1; 25,564 annotated genes). We combined predictions from the two references with the GeMoMa Annotation Filter (GAF), producing 18,231 gene models, and retained the 2,143 that did not overlap any TSEBRA locus (using the bedtools v2.29.2 subtract -A option) (*60*) as gap-filling models. We merged these with the TSEBRA annotation using AGAT v1.2.0 (agat_sp_merge_annotations.pl) (*61*), which yielded 24,668 protein-coding gene models after AGAT’s reconciliation of gene and transcript features.

#### UTR annotation with PASA

Only a minority of gene models in the merged annotation carried UTR annotations: BRAKER3 and GeMoMa predict coding sequence alone, so UTR features (4,540 5′ and 4,290 3′) belonged exclusively to the 2,880 models derived from full-length Iso-Seq transcripts, for which TransDecoder defines UTRs as the regions flanking the predicted ORF. To annotate UTRs comprehensively, we ran PASA v2.5.3 (*62*) with the 104,871 Iso-Seq transcripts supported by ≥5 full-length reads as evidence. We first cleaned transcripts with seqclean (*63*) (1,574 trimmed, none removed), then aligned them to the reference genome with GMAP v2021-12-17 (*64*) and clustered them into 52,515 PASA assemblies. We loaded the 24,668-gene annotation into the PASA database and compared it against these assemblies to identify transcripts supporting UTR extensions and gene-model refinements. A second round of comparison did not produce further updates, indicating convergence. PASA merged 131 sets of adjacent gene models supported by spanning Iso-Seq assemblies, absorbing 264 gene models into 131 merged models and reducing the total from 24,668 to 24,535. PASA added UTR annotations to 22,485 gene models (92%), increasing 5′ UTR features from 4,540 to 30,405 and 3′ UTR features from 4,290 to 25,320. The PASA-updated annotation comprised 24,535 protein-coding gene models and 37,655 transcripts.

#### Gene-model curation

Following PASA annotation, some genes were split across adjacent records sharing the same symbol. We reassigned gene symbols by reciprocal CDS overlap (≥0.30) with the reference annotation and merged records corresponding to a single reference gene, reducing the annotation from 24,535 to 22,488 genes (37,617 transcripts). We then removed spurious 1–2 bp UTR features introduced during format reconciliation with AGAT, along with 199 transcripts with a total CDS length shorter than 30 bp, arising from incomplete reference-gene projections, removing 162 genes. Of the remaining 22,326 genes, 379 (377 of them homology-based projections) had a representative CDS that could not be translated without internal stop codons, mostly due to frameshifts. For 62 of these, other isoforms encoded valid proteins. We removed the remaining 317, all single-transcript models. The final annotation comprised 22,009 genes and 37,101 transcripts, with 30,099 5′ and 25,218 3′ UTR features.

#### Annotation quality assessment

We assessed annotation completeness with BUSCO v5.5.0 (*65*) against the aves_odb10 lineage dataset (n = 8,338) in protein mode, using a single representative (longest) protein per gene extracted with AGAT (agat_sp_keep_longest_isoform.pl). Excluding 62 genes where the longest isoform had a disrupted reading frame, the 21,947 representative proteins yielded a BUSCO completeness score of 96.9% (96.4% single-copy, 0.5% duplicated), with 0.8% fragmented and 2.3% missing BUSCOs. The BUSCO dataset includes only conserved single-copy orthologs, most of which were present in the previous annotation, therefore newly recovered genes largely fall outside this set (and 81% of them lack a gene symbol).

### Molecular sex determination

Brown females were identified in the field, as most males were molting into reproductive plumage. One individual (D179028) showed brown plumage but patterns of gene expression resembling those of males, so we confirmed its sex by molecular sexing based on the CHD1 intron length polymorphism (*66*), alongside two known males and two known females as controls.

DNA was extracted from feather follicles after bead mill homogenization using the Qiagen DNeasy Blood & Tissue Kit following the manufacturer’s protocol. PCR amplification was performed in 10 μL reactions containing 1× reaction buffer, 4 mM MgCl₂, 0.25 mM each dNTP, 0.5 μM each primer, and 1 U Taq DNA Polymerase (Thermo Fisher, 10966-034), with 20– 100 ng template DNA. The forward primer P2 (5′-TCTGCATCGCTAAATCCTTT-3′) was labeled with FAM fluorescent dye and the reverse primer P8 (5′-CTCCCAAGGATGAGRAAYTG-3′) was unlabeled. Cycling conditions were 94 °C for 2 min; 35 cycles of 94 °C for 30 s, 49 °C for 45 s, 72 °C for 45 s; and a final extension at 72 °C for 5 min. Fragment sizes were determined by running FAM-labeled PCR products on a Fragment Analyzer alongside an internal size standard, with males (ZZ) expected to yield a single *CHD1-Z* fragment of ∼347 bp and females (ZW) two fragments (∼347 bp *CHD1-Z* and ∼385 bp *CHD1-W*). D179028 yielded a single fragment, confirming the sample came from a male. All sex assignments are reported in Table S1.

### Differential expression analysis

Throughout, we refer to candidate genes (*ASIP*, *TYRP1*, *OCA2*, *SLC45A2*, *HERC2*, *MLANA*, *KITLG*, *GPT2*, *MYO5A*, *DCT* and *CAMK2D*), defined as genes from the differentiated genomic regions identified by previous population-genomic scans of the radiation (*12*, *15*). In the most recent of these scans (*15*), F_ST_ was estimated per site and in 10 kb windows for 15 pairwise comparisons among six species. Outlier windows were those associated with plumage color in a GWAS or in the top 0.1% of weighted F_ST_ in any of the 15 pairwise comparisons and containing at least one variant with F_ST_ > 0.75, and consecutive outlier windows were merged into divergence peaks. The analyses described below as candidate-gene analyses are restricted to genes within the divergence peaks, and composed primarily but not exclusively of melanogenesis genes (*GPT2* and *HERC2* are not involved in melanogenesis).

#### Sequencing read quantification

We quantified gene-level read counts from the aligned BAM files with featureCounts v2.0.3 (subread v2.1.1) (*67*), using the annotation prior to the final curation step (22,326 genes; see Gene-model curation) converted to GTF format with gffread v0.9.12 (*68*), giving a matrix of 22,326 genes across 46 samples. Reads were counted at the gene level over exon features (-t exon -g gene_id), with paired-end counting (-p --countReadPairs), reverse-strand specificity (-s 2), and a minimum mapping quality of 10 (-Q 10). These parameters produced an average assignment rate of 66.6%. We then excluded the 317 genes removed during curation, giving a matrix of 22,009 genes across 46 samples.

#### Sample quality control

We assessed quality across all 46 samples using three complementary quality-control criteria: assigned read pairs relative to the median, distance from the library’s patch centroid on the first two axes of a global principal component analysis (PCA) of variance-stabilized counts (VST), and mean Pearson correlation with all other libraries based on the same transformed counts. A library was flagged on a criterion if its assigned reads fell below 0.5× the median, its distance from the patch centroid exceeded the median by more than three median absolute deviations (MAD), or its mean correlation fell below the median by more than three MAD. Samples flagged on two or more criteria were excluded.

Across the 46 samples, assigned read pairs ranged from 18.6 to 71.2 million (median 51.0 million), and mean inter-sample correlation ranged from 0.93 to 0.98 (mean 0.97). Three samples failed two or more criteria (D165854_SP_CR_BR, C53148_CIN_BR_CH, and C65600_PIL_CR_BC) and were excluded, leaving 43 samples from 29 individuals. All patch-color combinations remained represented after exclusion. Per-sample quality metrics are shown in Fig. S6.

#### Filtering, normalization, and transformation

Genes were retained if they reached at least 10 counts in three or more samples, leaving 16,811 genes. All differential-expression analyses were performed with DESeq2 v1.50.2 (*69*) in R v4.5.1 (*70*). For all ordinations and per-gene visualizations we used a single variance-stabilizing transformation (VST) computed with an intercept-only design (∼ 1) and the blind option, so that experimental factors did not inform the transformation. The same blind VST matrix underlies every PCA, gene panel, and heatmap, ensuring a consistent set of transformed values throughout.

#### Differential gene expression: significance criteria and filtering

Throughout, the focal term is written last in each model design formula. The term order does not affect the fit. We used one set of significance criteria across all models. A gene was considered differentially expressed in a contrast if it reached a Benjamini–Hochberg adjusted *p* < 0.05, an absolute log2 fold-change ≥ 1, and a mean normalized expression (baseMean) ≥ 100. Log2 fold-changes were shrunk for all pairwise contrasts, using apeglm (*71*) for the two-level sex contrast and the adaptive shrinkage estimator ashr (*72*) for multi-level pairwise contrasts, as apeglm supports only contrasts against the reference level whereas ashr accommodates arbitrary pairwise contrasts with comparable empirical-Bayes shrinkage.

Two filters were applied to avoid inflated fold-changes from genes expressed in one group but absent from another. First, a presence filter required each gene to be expressed (≥ 10 counts in ≥ 2 samples) in at least two levels of the focal variable. Genes that met the expression threshold in only one group, or in neither group, were excluded from fold-change-based tests because fold changes are not reliably estimated when a gene has negligible or undetectable expression in one group. Second, contrasts with an absolute shrunken log2 fold-change above 10 were excluded as extreme fold-change estimates. However, no contrast exceeded this threshold after shrinkage. A focal level represented by fewer than three libraries was not tested, though the corresponding samples were retained in ordinations and heatmaps so that every sample appears.

For analyses comprising several pairwise contrasts, each gene was annotated with the number of contrasts in which it was significant, and genes significant in two or more were flagged as robustly differentially expressed. Gene sets underlying ordinations and heatmaps were the union of genes significant in any pairwise contrast within a model, restricted to gene models with an assigned gene symbol, ordered by number of significant contrasts and then adjusted *p*-value. All differential-expression tests were Wald contrasts.

#### Differential gene expression between feather patches

Differential expression among feather patches (crown, breast, back) used a primary model of ∼ sex + color + patch fitted to all 43 samples, so that patch effects are estimated while controlling for sex and color. Pairwise contrasts (crown vs. breast, crown vs. back, breast vs. back) were extracted with Wald tests and ashr shrinkage. We compared the primary model against three alternatives, each removing a different potential artifact: an uncontrolled model (∼ patch), which shows the raw patch difference before adjustment; a males-only model (∼ color + patch), which eliminates sex; and a within-individual paired model (∼ individual + patch), restricted to the 11 birds sampled at more than one patch, which removes the non-independence arising from repeated sampling of the same individual. The paired model cannot separate patch from color, since color varies with patch within an individual. The males-only model retains color in the design but treats libraries from the same bird as independent. These models therefore address complementary limitations. Because patch and color are partly collinear, their effects can only be separated for colors sampled across multiple patches.

#### Color-associated gene expression

Plumage color varies both within and between feather patches, but the two are non-independent: each patch contains a different, unbalanced subset of colors, and some colors occur in only one patch (Fig. 1A). Color is also confounded with species: in our sampling each color category was dominated by one or two species (e.g. salmon was exclusively *S. pileata*, gray almost entirely *S. melanogaster*; Fig. 1A). Models cannot therefore estimate a color effect independent from species. We therefore compared color-associated expression across different models to assess how much of any color signal is separable from patch.

Our primary color analyses were run in males, removing the sex-confounded brown females, and restricted to color categories represented by at least three samples, which excludes brown, leaving 34 samples. We counted color-associated genes (pairwise Wald contrasts, criteria as above) under two designs: patch-controlled (∼ patch + color, the primary model for color), in which colors are compared after adjusting for body region, and uncontrolled (∼ color), which retains any difference in patch composition between color groups. To test brown, which is absent from the male subset, we additionally fitted all 43 samples with a sex term (∼ sex + patch + color).

We also tested color separately within each patch in which more than one color was testable (crown: black, gray; breast: black, salmon, orange; back: salmon, gray), using pairwise Wald contrasts. Brown crowns were not testable in the male-only subset because only one male brown crown was sampled, below the minimum of three samples required per color group. We therefore also tested crown color across all samples using a model including sex as a covariate (∼ sex + color).

#### Sex-biased gene expression

Sex-biased expression was identified with a primary model of ∼ sex (reference level: male) fitted to all 43 samples, restricted to the 15,760 genes reaching at least 10 counts in at least two samples of both sexes. The female-versus-male contrast was tested with a Wald test and its log2 fold-change shrunk with apeglm (*71*). Negative values therefore indicate higher expression in males. We assessed robustness against two further models additionally controlling for patch (∼ patch + sex) and for patch and color (∼ patch + color + sex), recording for each gene the number of models in which it was significant.

Because sex is constant within an individual, repeated samples from different patches of the same bird carry no additional information about sex, and the sample size is the number of individuals (24 males, five females) rather than the number of samples. To assess the influence of this non-independence, we refitted the sex model using one randomly chosen sample per individual, repeating the procedure 20 times and recording for each gene the proportion of draws in which it met the significance criteria. We ran this subsampling under two designs, ∼ sex and ∼ patch + sex. Because each draw includes a single patch per bird, differences among patches add noise to the unadjusted model and can hide a sex effect in genes with expression varying strongly among patches. The patch-adjusted design accounts for this variation.

#### Sex-chromosome linkage and dosage

Birds are only partially dosage-compensated (*73*, *74*), so Z-linked genes are expected to be male-biased in proportion to gene copy number. A completely uncompensated Z-linked gene yields a female-versus-male log2 fold-change of −1. Because the Z chromosome is split across multiple contigs, we classified genes as Z-linked by symbol match to the Zebra Finch Z gene set (n = 1,219) rather than by chromosome assignment. Symbol matching alone was insufficient, as nomenclature differs between annotation versions. We therefore also identified Z-fragment contigs empirically as those on which at least 50% of tested genes (minimum five) matched the Zebra Finch Z chromosome, and classified all genes on them as Z-linked. Six contigs met this criterion (7, 28, 35, 41, 52, 57). Genes that matched the Zebra Finch Z gene set and did not fall on a Z-fragment contig were treated as autosomal. We tested enrichment of Z-linked genes among sex-biased genes with Fisher’s exact test. To characterize dosage compensation we computed, for each Z-linked gene, the ratio of mean female to mean male expression directly from size-factor-normalized counts rather than from shrunken coefficients, which bias fold-changes toward zero and would understate incomplete compensation. A female-to-male ratio near 0.5 is the null expectation for an uncompensated Z-linked gene, so we interpreted sex bias at Z-linked genes against this expectation (that is, as the degree of dosage compensation) rather than as evidence of sex-specific regulation, and report them separately from autosomal genes.

### Differential transcript usage

Transcripts were quantified with Salmon v1.10.0 (*75*) against an index that includes the whole genome alongside the transcriptome so that reads from unannotated regions are not forced onto transcripts. Sequence and GC-bias correction were enabled (--seqBias --gcBias -- validateMappings). Quantifications were imported with tximport v1.38.2 (*76*) as length-corrected counts (scaledTPM), so that isoform values reflect relative usage rather than transcript length. Analyses were restricted to candidate pigmentation genes with at least two expressed isoforms.

Differential transcript usage (DTU) was tested with DEXSeq v1.56.0 (*77*), applied at the transcript level by treating each gene’s transcripts as the units among which usage is compared. The model has three terms. The sample term accounts for differences in sample-level abundance, while the exon term captures baseline differences in isoform usage. The condition × exon interaction tests whether the usage of a given transcript, relative to the other transcripts of the same gene, differs between the groups being compared. A significant interaction therefore means that the gene’s isoform composition changes between groups, regardless of whether the gene’s total expression changes. The model is fitted as ∼ sample + exon + condition:exon, and significance is assessed per transcript. Per-gene significance was summarized as the minimum Benjamini–Hochberg adjusted *p*-value across a gene’s transcripts.

For each test we recorded the largest change in any isoform’s proportion between groups and, separately, the change in the proportion of the dominant (highest mean expression) isoform, together with the group levels at which that proportion was lowest and highest. Because DEXSeq is sensitive at very small effect sizes, several tests reached adjusted *p* < 0.05 with small dominant-isoform shifts. We therefore report genes as showing differential transcript usage at adjusted *p* < 0.001, and give adjusted *p*-values under both thresholds in Table S10.

### Individual genotyping at candidate variants and species clustering

To evaluate whether individuals cluster by species based on the outlier variants identified from the population-genomic scans when genotyped directly from RNA-seq reads, we performed PCA. Reads from different libraries from the same individual were pooled before genotyping.

Per-patch BAM files were therefore merged with *samtools* (v1.20; (*78*)) under a single individual identifier, allowing sequence coverage to accumulate across patches so that each individual contributed a single genotype at each site. Individuals for which the only available library failed quality control (D165854 and C53148) were excluded from the genotype dataset.

Genotypes were called across individuals using *bcftools mpileup* and *bcftools call* (v1.20; (*78*)) under the multiallelic calling model (-m). Heterozygous genotypes were retained, as RNA-seq libraries can contain transcripts from both alleles. We considered three variant sets: outlier variants located within candidate pigmentation genes, all F_ST_/GWAS outlier variants (*15*), and variants within the 500 most highly expressed genes. The top-500 genes were ranked by mean variance-stabilized expression, and variants overlapping candidate pigmentation genes were removed from this background set so that it represented a candidate-independent set of highly expressed genes.

A single position can fall in different features of different isoforms or genes (for example CDS in one isoform and intergenic in another). Each site was therefore assigned to a single class in the priority order CDS, UTR, intron, and finally intergenic. For the PCA analyses we retained biallelic SNPs with QUAL ≥ 30 classified as CDS or UTR variants. PCA analyses were conducted separately for each set of variants and using the combined CDS+UTR set, the CDS-only, or the UTR-only subsets.

A genotype was considered reliable only when the combined read depth across an individual’s multiple samples was at least five. Genotypes below this threshold were set to missing. Sites genotyped in fewer than 70% of individuals were excluded, and remaining missing genotypes were set to the mean reference-allele dosage across all individuals genotyped at that site, since PCA requires a complete matrix. Because imputed values equal the site mean, imputed individuals do not contribute signal at that site after centering. Genotypes were coded as reference-allele dosage (0, 1, or 2) and standardized per site, and sites left invariant by filtering were removed before PCA. Across all analyses, PC1 was uncorrelated with per-individual missingness (|r| ≤ 0.16 for the candidate-gene sets), indicating that clustering reflects genotype rather than coverage.

The final genotype dataset comprised 35 individuals, the 29 from the expression analyses and six pilot individuals (Table S1), yielding 47 candidate-gene sites (15 CDS, 32 UTR), 345 all-outlier sites (191 CDS, 154 UTR), and 50,736 variable (non-outlier) CDS+UTR sites in the candidate-excluded top-500 set (16,089 CDS and 34,647 UTR). Candidate positions not recovered were predominantly UTR variants in transcripts with low expression in feather follicles, where coverage is insufficient for a genotype call. Results were unchanged when restricted to the 29 individuals in the expression analyses.

### Consequence prediction of candidate outlier variants

We extracted all outlier positions falling within the annotated span of each of eight candidate pigmentation genes from the F_ST_ and GWAS outlier VCF (*15*), retaining all variant classes, and ran *bcftools csq* (*79*) with the improved annotation and the *S. hypoxantha* reference genome. Because genotypes were unphased, we ran csq in local mode (-l), evaluating each variant independently rather than combining nearby variants into haplotypes. Unphased heterozygous genotypes were merged (-p m) and the effect of mutations was predicted for all samples (-s -). Transcript identifiers returned by csq were mapped to gene symbols using the transcript ID and gene name attributes of the annotation. This yielded 591 positions in eight candidate genes. Under the improved annotation, three positions within these candidate-gene intervals fell in transcripts of adjacent non-candidate genes and were excluded, leaving 588 positions in eight candidate genes.

Because amino-acid coordinates differ among isoforms, variants leading to protein changes are reported against the isoform with the highest mean expression across libraries (Salmon transcript-level TPM) among those predicted to carry the variant. Alternative isoforms carry substitutions at different residues: the *SLC45A2* change at contig 7:5,261,935 is Asn368Asp on the dominant isoform (55% of gene expression) and Asn350Asp or Asn341Asp on minor isoforms, and the *OCA2* change at contig10:6,277,903 is Met146Thr on two minor isoforms (15% combined) and Met126Thr on the four isoforms carrying 85% of expression.

For the candidate-gene outlier positions in coding and untranslated sequence, we estimated alternate allele frequencies in each species from the multi-sample VCF of F_ST_ and GWAS outliers (*15*), which comprises 10,762 positions genotyped in 127 individuals from ten species of the radiation. Because per-site coverage was low (median depth 5), frequencies were computed from pooled reference and alternate read counts across the individuals of each species rather than from individual genotype calls. At multiallelic positions, all non-reference read counts were pooled. Frequencies computed instead from genotype calls (minimum depth 5) gave similar values across the candidate positions (median absolute difference 0.02 across 1,062 species-by-site estimates), with the largest discrepancies confined to species represented by fewer than four genotyped individuals.

### Analysis of feather carotenoid content

We used high-performance liquid chromatography with ultraviolet and visible light detection (HPLC-UV-Vis) to characterize and quantify feather carotenoid content, following established methods (*80*). We trimmed distal barbs from three feathers from each patch (∼0.9 mg) into a pre-weighed 2 mL screwcap centrifuge tube. Carotenoid analyses were performed on a set of 15 individuals (29 patch samples) partly overlapping the RNA-seq species set (Table S8). Feathers for these analyses were sampled from wild-caught adult individuals as part of a previous study (*35*), and include three species not represented in the RNA-seq dataset: *S. hypochroma* (Rufous-rumped Seedeater), *S. iberaensis* (Iberá Seedeater), and *S. ruficollis* (Dark-throated Seedeater). We weighed to the nearest 0.01 mg the tube containing the feather sample with an analytical balance, and then calculated sample mass as the difference between the measures. We then added 1 mL of methanol (Fisher Scientific Inc.) and 0.1 g of zirconium oxide beads (Zrob20, Next Advance, Inc.) and homogenized at 4 kHz for 8 min. on a beadmill (BeadBug, Benchmark Scientific Inc.). We then centrifuged the sample at 15,000 g and collected the supernatant in a 15 mL conical tube. We repeated this methanol extraction twice more, combining the supernatants from each repetition. This was followed by two extractions with 1 mL each of hexane:tert-methyl butyl ether (1:1, vol:vol, Fisher Scientific Inc.) with the same grinding and separation process. After five rounds of extractions were completed, we washed the supernatant by adding 3 mL of 0.9% NaCl in distilled deionized water, mixed by vortexing for 30 sec, and centrifuging the sample at 2,000 rcf for 5 min. We then transferred the upper solvent fraction to a 2 mL glass vial and dried under a stream of nitrogen gas. For HPLC analysis, we dissolved the dried extract in 120 µL of mobile phase solvent (described below) and injected 100 µL of the sample into an Agilent 1260 HPLC fitted with a C30 carotenoid column (5.0 μm, 4.6 × 250 mm, YMC, CT99S05-2546WT). We separated the carotenoids with a mobile phase of methanol:acetonitrile:tetrahydrofuran (45:45:10; vol:vol:vol) through 11 mins, a gradient up to 35:35:33 through minute 21, then isocratic at 35:35:33 through 24 minutes, and then a return to 45:45:10 through 30 mins. The flow rate was constant at 1.2 mL min-1 and the column temperature maintained at 30℃. We monitored samples with a UV-Vis photodiode array detector and identified carotenoids by comparing their retention times and UV-Vis spectra to external zeaxanthin, lutein, and β-carotene (dsm-firmenech, Stroe, Netherlands). Carotenoid concentrations were quantified based on an external standard curve for zeaxanthin.

## Acknowledgments

Sequencing was performed by Transcriptional Regulation and Expression (TREx) Facility at the Cornell University Biotechnology Resource Center (RRID:SCR_022532) and Weill Cornell Medicine Genomics Resources Core Facility (RRID:SCR_025634). Computational analyses were performed using the Cornell University BRC Bioinformatics Core Facility (BioHPC, RRID:SCR_021757). We are grateful to the Centro Nacional de Pesquisa e Conservação de Aves Silvestres (CEMAVE, Brazil; permit: 5050/1) and the Instituto Chico Mendes de Conservação da Biodiversidade (ICMBio, Brazil; SISBIO permit: 95456). We thank the Uy lab at the University of Rochester for valuable comments on the manuscript.

## Funding

This study was funded through a grant from the National Science Foundation (USA, DEB - 2232929).

## Author contributions

Conceptualization: M.Rec., L.C.

Methodology: M.Rec., M. Rep., L.C.

Investigation: M.Rec., L.C., S.K., J.R.R.R., M.B.T, L.F.S., C.K.

Formal analysis: M.Rec.

Resources: M.Rep., M.B.T., L.F.S., C.K., L.C.

Data curation: M.Rec., M. Rep.

Visualization: M.Rec.

Funding acquisition: L.C.

Supervision: L.C.

Project administration: L.C.

Writing—original draft: M.Rec., L.C.

Writing—review & editing: M.Rec., M.Rep., M.B.T., S.K., J.R.R.R., L.F.S., C.K., L.C.

## Competing interests

The authors declare that they have no competing interests.

## Data, code, and materials availability

Raw sequencing data have been deposited in the NCBI Sequence Read Archive under BioProject PRJNA1523881 and will be publicly released upon publication. PacBio Kinnex Iso-Seq reads are under accession SRR40711656, and the 57 Illumina RNA-seq libraries under accessions SRR40711640 to SRR40711655 and SRR40711657 to SRR40711697; per-library accessions are given in Table S1. The improved *Sporophila hypoxantha* genome annotation (GFF3 and GTF files), the transcript-to-gene table, the gene-level count matrix, sample metadata, transcript-level Salmon quantifications, and the code for the differential expression, differential transcript usage, variant classification, and genotype PCA analyses are available on Figshare (https://doi.org/10.6084/m9.figshare.34862871).

## Notes

### Competing Interest Statement

The authors have declared no competing interest.

