## Supplementary Material for "Plumage diversity without differential expression of pigmentation genes in a rapid avian radiation"

**This PDF file includes:**

Figs. S1 to S6
Tables S1 to S10

Contents

### Table of Contents

*Supplementary Figures* ... *3*

**Figure S1. Differentially expressed genes among feathers from different body patches** ... **4**

**Figure S2. Z-chromosome dosage compensation and autosomal sex-biased gene expression** ... **5**

**Figure S3. Reannotation of the outlier variants from previous GWAS and $F_{ST}$ scans** ... **7**

**Figure S4. Differential transcript usage (DTU) at candidate pigmentation genes.** ... **8**

**Figure S5. Principal component analyses (PCA) of RNA-seq-derived genotypes in 35 individuals for three variant sets.** ... **9**

**Figure S6. Quality-control assessment of the 46 RNA-seq samples** ... **10**

*Supplementary Tables* ... *11*

**Table S1. Sample metadata and sequencing summary** ... **11**

**Table S2. Number of differentially expressed genes in each pairwise patch contrast across four experimental designs** ... **14**

**Table S3. Differentially expressed genes across the three pairwise patch contrasts under the primary model** ... **15**

**Table S4. Number of differentially expressed genes in each pairwise color contrast across seven experimental designs** ... **20**

**Table S5. Differentially expressed genes in the color contrasts** ... **21**

**Table S6. Number of sex-biased differentially expressed genes** ... **23**

**Table S7. Sex-biased differentially expressed genes** ... **24**

**Table S8. Feather carotenoid content by individual** ... **27**

**Table S9. Predicted consequences and species allele frequencies of outlier variants within candidate pigmentation genes** ... **29**

**Table S10. Differential transcript usage at candidate pigmentation genes** ... **32**

#### Supplementary Figures

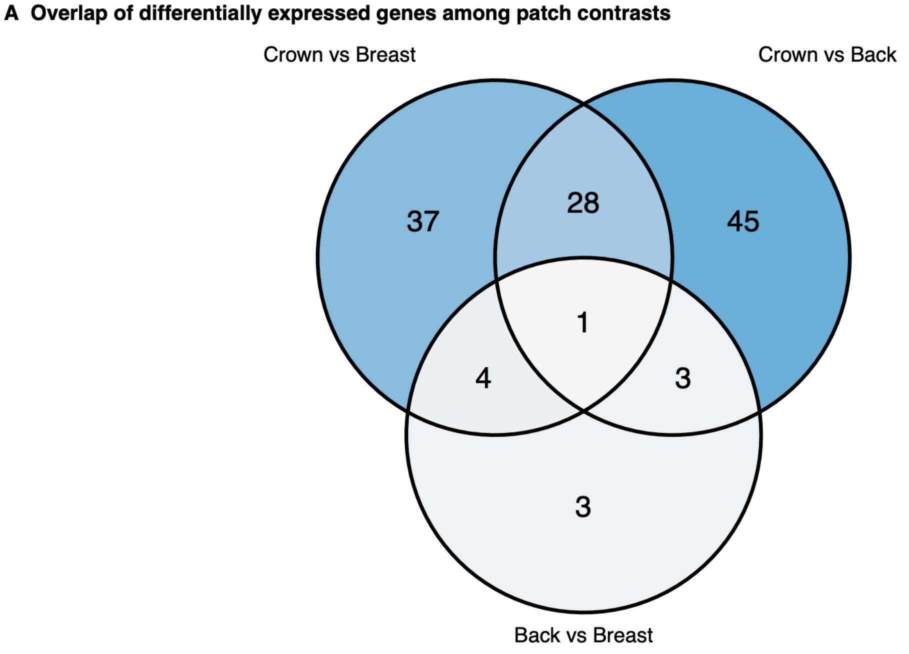

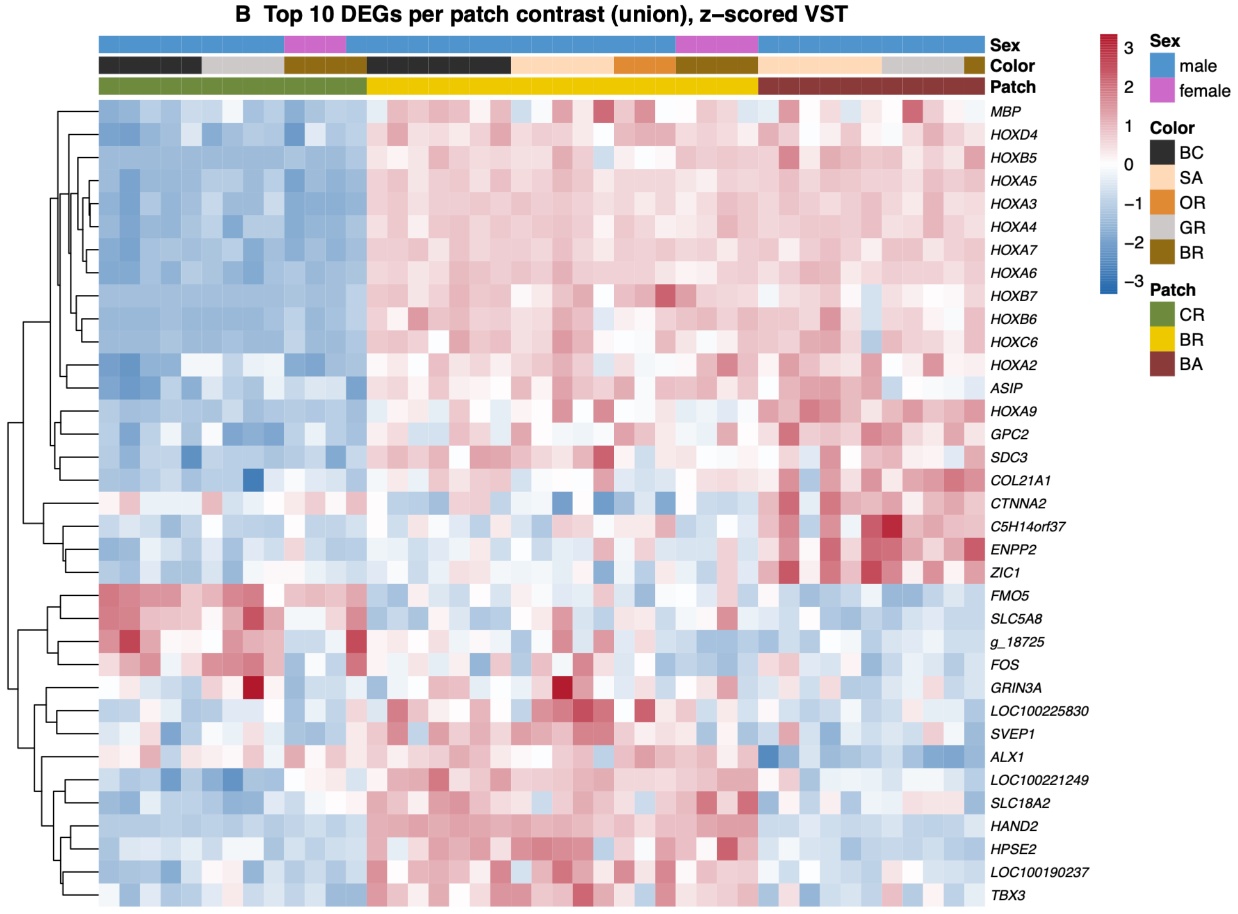

**Figure S1. Differentially expressed genes among feathers from different body patches. (A)** Three-way Venn diagram of genes differentially expressed in each pairwise patch contrast under the primary model (~ sex + color + patch). Of 121 differentially expressed genes, most were specific to a crown contrast (37 for crown vs. breast, 45 for crown vs. back), with fewer distinguishing breast from back (3). Only *HAND2* was differentially expressed in all three contrasts. **(B)** Heatmap of the 35 genes resulting from the union of the top 20 genes (by adjusted *p*) from each pairwise contrast (only 11 significant genes included for breast vs back). Values are per-gene z-scores of the blind (intercept-only) variance-stabilized counts. Columns are the 43 samples, ordered by patch (crown, breast, back) then plumage color, with patch, color, and sex annotated above. Fold-change and significance are from Wald tests with ashr shrinkage.

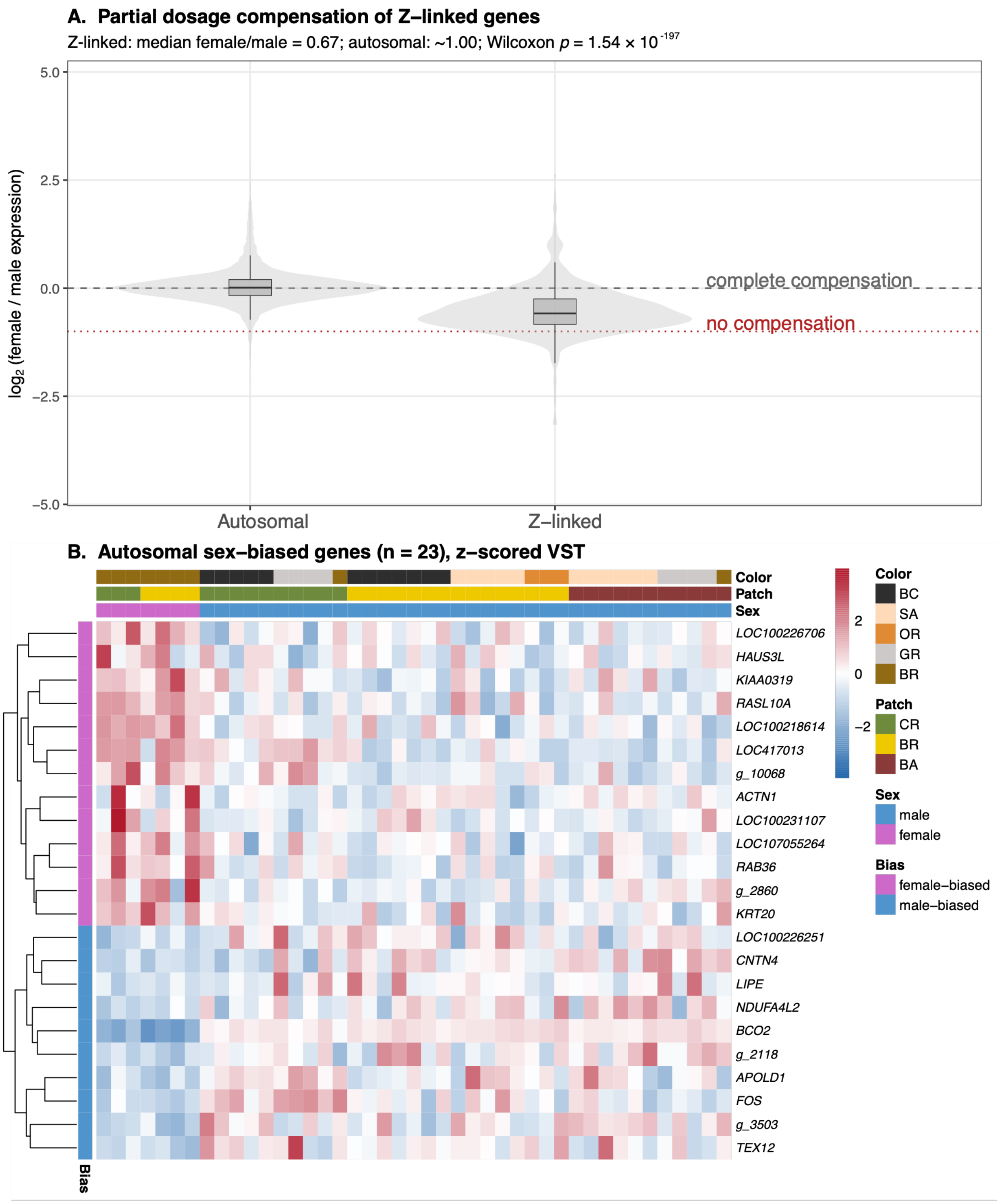

**Figure S2. Z-chromosome dosage compensation and autosomal sex-biased gene expression. (A)** Distribution of the female-to-male expression ratio (log2, computed from size-factor-normalized counts) across all tested genes, separated into autosomal and Z-linked classes. Violins show the distribution and boxplots the median and interquartile range. The dashed line marks complete dosage compensation (equal female and male expression, log2 ratio 0) and the dotted red line marks no compensation (females show half the level of male expression, log2 ratio −1). Z-linked genes have a median female-to-male ratio of 0.67, intermediate between the two lines, indicating an overall partial dosage compensation of the expression of Z-linked genes in our dataset (Wilcoxon rank-sum test, autosomal vs Z-linked, $p = 1.54 \times 10^{-197}$). Genes were assigned to classes by symbol match to the Zebra Finch Z gene set and by empirically identified Z-fragment contigs (Methods). **(B)** Heatmap of the 23 autosomal genes with sex-biased expression, showing per-gene z-scores of the blind variance-stabilized counts. Columns are the 43 samples ordered by sex (females first) and then patch. Sex, patch, and plumage color are annotated above the heatmap and the direction of the sex bias (higher male or female expression) is annotated on the left. Rows represent the expression level of different genes.

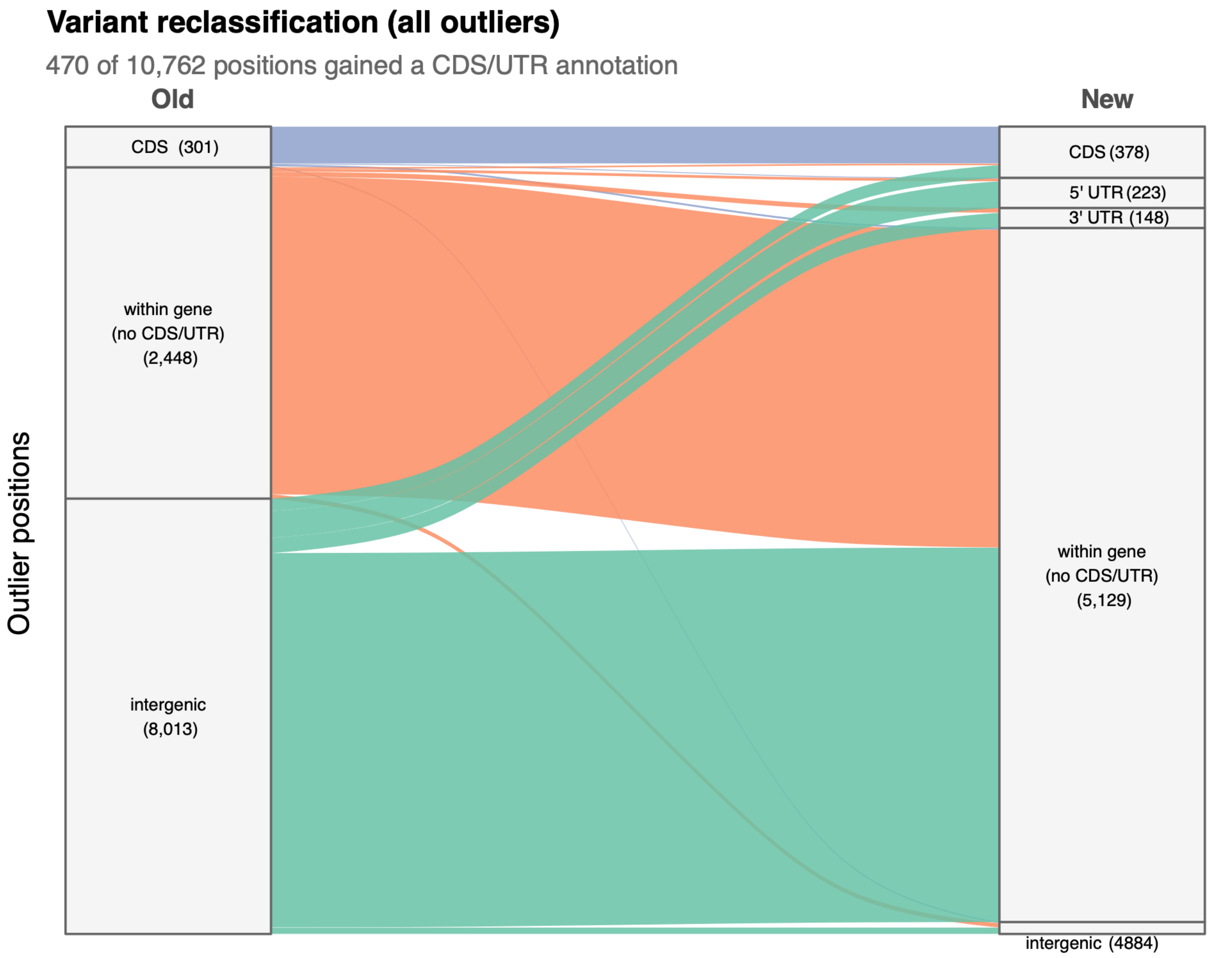

**Figure S3. Reannotation of the outlier variants from previous GWAS and $F_{ST}$ scans.** Reclassification of all outlier positions according to the improved genome annotations. Of 10,762 outlier positions, 470 gained a CDS or UTR annotation following reannotation. The flow between intergenic variants is represented in a 1:100 scale for ease of visualization.

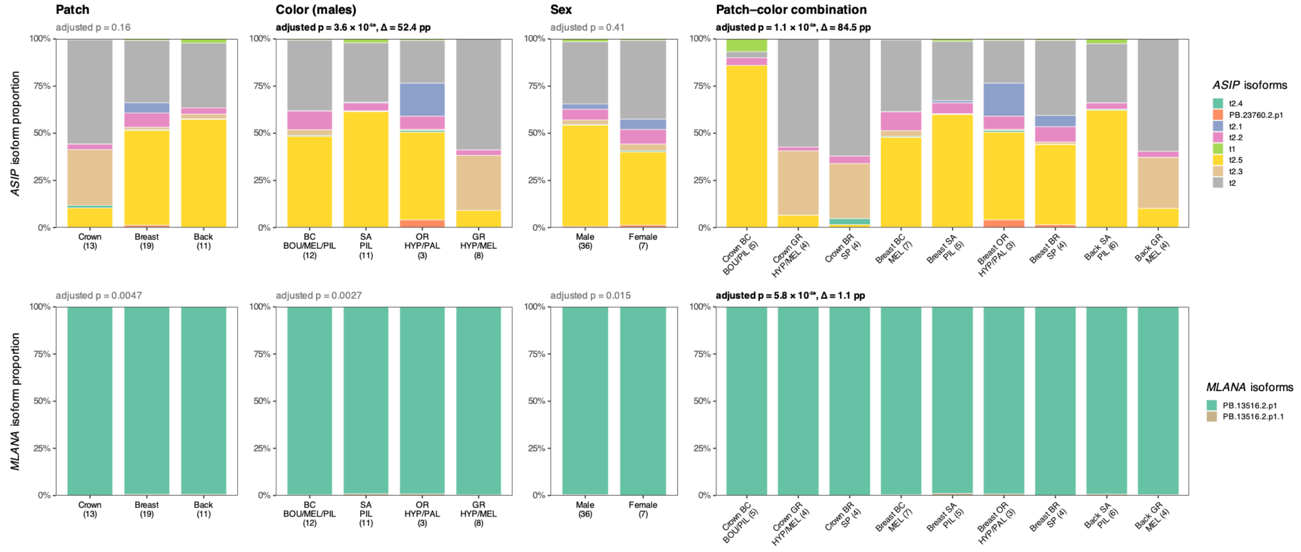

**Figure S4. Differential transcript usage (DTU) at candidate pigmentation genes.** Each row shows one gene, and each column one of the four groupings tested in Fig. 4B: feather patch, plumage color (males only), sex, and patch–color combination. Stacked bars give the proportion of the gene's total expression contributed by each isoform (colors identified at right), calculated from scaled TPM counts pooled across the samples in each group. Groups are labelled below each bar with the species they contain and, in parentheses, the number of samples. The adjusted *p*-value for each gene and grouping is shown above the panel, as in Fig. 4B. Values with adjusted $p < 0.001$ are in bold and marked with an asterisk, followed by Δ value, which is the difference in percentage points (pp) in the proportion of the gene's expression contributed by its dominant isoform between the groups where it is most and least abundant. In *MLANA* the second isoform accounts for less than 2% of expression in every group, so the bars appear uniform. *ASIP* is expressed at low levels in crown feathers (221 to 1,536 counts per patch–color group, against 3,245 to 33,732 in the breast and back), so isoform proportions in the three crown groups are calculated based on comparatively fewer reads. Color codes: BC, black; SA, salmon; OR, orange; GR, gray; BR, brown. Species codes: BOU, *S. bouvreuil*; HYP, *S. hypoxantha*; MEL, *S. melanogaster*; PAL, *S. palustris*; PIL, *S. pileata*; SP, brown individuals not assignable to species by plumage.

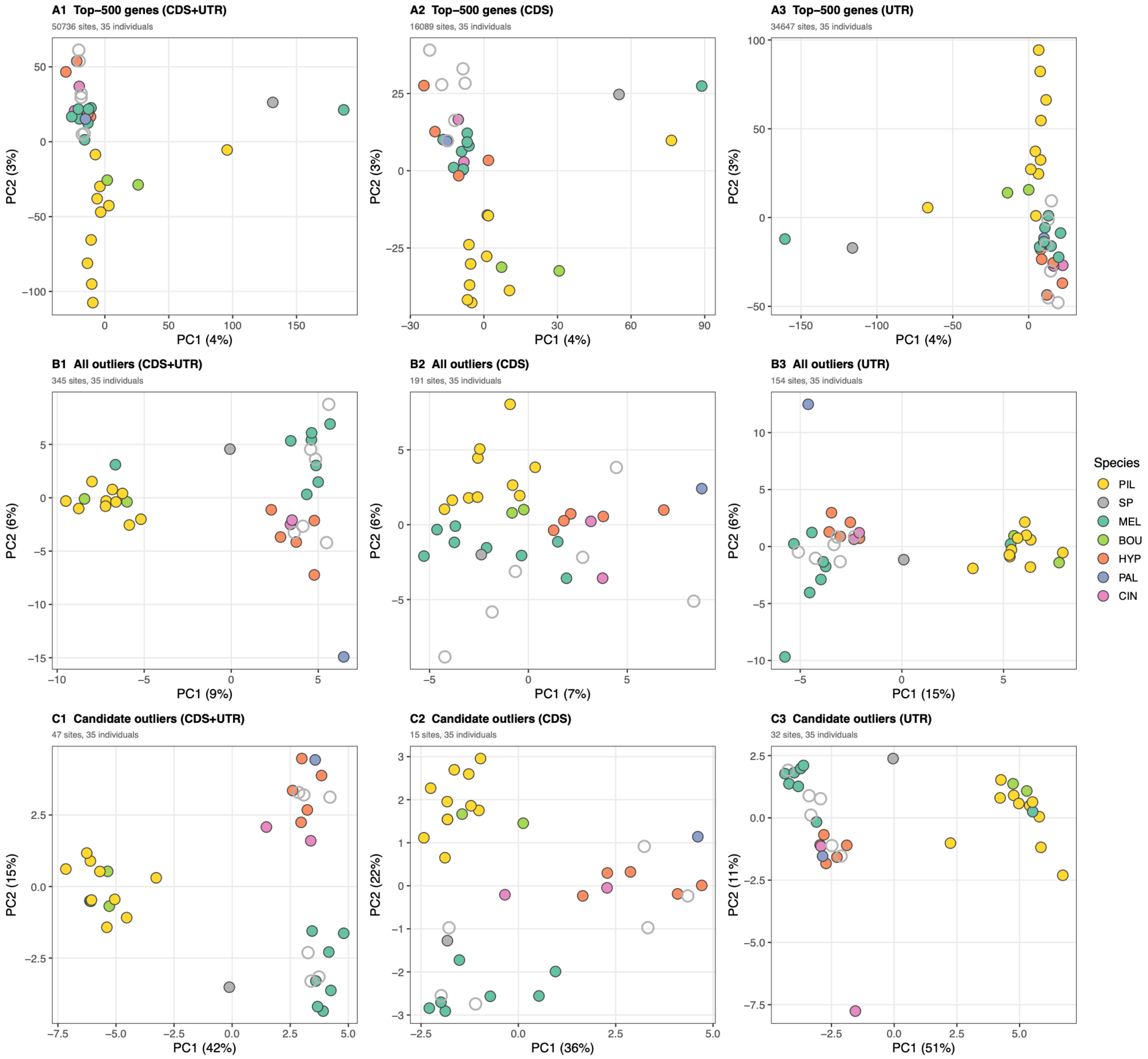

**Figure S5. Principal component analyses (PCA) of RNA-seq-derived genotypes in 35 individuals for three variant sets.** Variants within the 500 most highly expressed genes, excluding candidate genes (**A1–A3**); all $F_{ST}$/GWAS outlier variants (**B1–B3**); and outlier variants within candidate genes (**C1–C3**). Left, combined CDS and UTR variants; center, CDS variants only; right, UTR variants only. The 35 individuals comprise the 29 used in the expression analyses and six from the pilot study. Genotypes were called from pooled RNA-seq samples per individual. Points are colored by species and open circles indicate brown individuals that cannot be assigned to species by plumage (females or unknown sex), and the filled gray circle is the brown male (D179028).

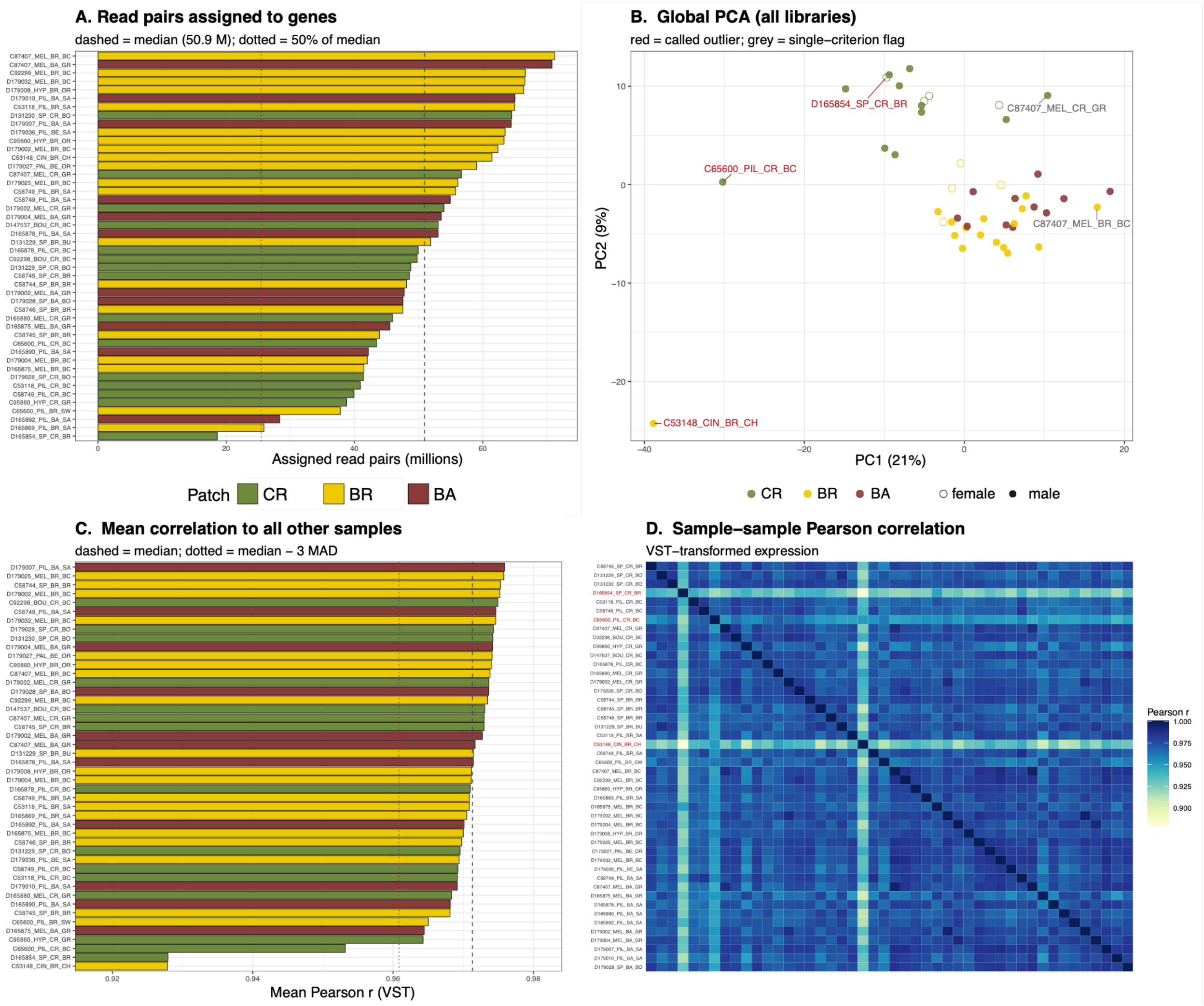

**Figure S6. Quality-control assessment of the 46 RNA-seq samples.** (**A**) Number of reads successfully assigned to genes for each sample. The dashed line indicates the median assigned-read depth (51 million reads), and the dotted red line indicates the threshold used to flag samples with fewer than 50% of the median assigned reads. (**B**) Principal component analysis (PCA) of variance-stabilized gene expression counts across all samples. Points are colored by sampling patch and shaped by sex. The labels identify samples flagged by the QC procedure. (**C**) Mean Pearson correlation of each sample with all other samples based on variance-stabilized expression. The dashed line indicates the median correlation and the dotted red line indicates the threshold defined as the median minus three median absolute deviations (MAD). (**D**) Pairwise Pearson correlations among samples based on variance-stabilized expression counts. Samples flagged by the QC procedure are indicated in red. Samples were considered outliers when they failed at least two independent QC criteria: low read depth, excessive distance from the patch centroid in PCA space, or low mean inter-sample correlation. Samples D165854_SP_CR_BR, C53148_CIN_BR_CH, and C65600_PIL_CR_BC were therefore excluded from downstream analyses.

#### Supplementary Tables

**Table S1. Sample metadata and sequencing summary.** Asterisks mark the three samples flagged on two or more quality-control criteria and excluded from downstream analyses, leaving 43 samples from 29 individuals. Per-sample statistics are shown in Figure S6. Patch codes: CR, crown; BR, breast; BA, back. Color codes: BC, black; SA, salmon; OR, orange; GR, gray; BR, brown. Sex: M, male; F, female; UNK, unknown. Samples marked with (#) were pooled for Iso-seq sequencing. Samples below the line are from the pilot study that were not used in the differential expression analysis. UNK, sex not determined (pilot samples used only for genotyping).

| Sample | NCBI SRA accession | Species | Patch | Color | Sex | Total reads | Uniquely mapped (%) | Feature counts assign. (%) | Read pairs assigned to features (Million) |
|---|---|---|---|---|---|---|---|---|---|
| C53118_PIL_BR_SA# | SRR40711697 | *S. pileata* | BR | SA | M | 99,464,324 | 80.6 | 66.7 | 65.0 |
| C53118_PIL_CR_BC | SRR40711696 | *S. pileata* | CR | BC | M | 67,606,375 | 73.0 | 67.7 | 41.0 |
| C53148_CIN_BR_CH* | SRR40711685 | *S. cinnamomea* | BR | OR | M | 94,190,273 | 76.9 | 63.3 | 61.6 |
| C58744_SP_BR_BR | SRR40711674 | *Sporophila sp.* | BR | BR | F | 99,798,213 | 60.6 | 64.9 | 48.2 |
| C58745_SP_BR_BR | SRR40711663 | *Sporophila sp.* | BR | BR | F | 79,490,846 | 72.8 | 59.9 | 43.9 |
| C58745_SP_CR_BR# | SRR40711652 | *Sporophila sp.* | CR | BR | F | 81,921,920 | 73.2 | 67.7 | 48.6 |
| C58746_SP_BR_BR | SRR40711643 | *Sporophila sp.* | BR | BR | F | 86,669,412 | 72.0 | 61.6 | 47.6 |
| C58749_PIL_BA_SA | SRR40711642 | *S. pileata* | BA | SA | M | 87,537,056 | 75.5 | 69.3 | 55.0 |
| C58749_PIL_BR_SA | SRR40711641 | *S. pileata* | BR | SA | M | 87,710,945 | 80.9 | 64.1 | 55.8 |
| C58749_PIL_CR_BC | SRR40711640 | *S. pileata* | CR | BC | M | 86,933,036 | 55.0 | 70.2 | 40.0 |
| C65600_PIL_BR_SW | SRR40711695 | *S. pileata* | BR | SA | M | 69,075,198 | 70.3 | 61.0 | 37.8 |
| C65600_PIL_CR_BC* | SRR40711694 | *S. pileata* | CR | BC | M | 81,833,096 | 64.0 | 68.5 | 43.6 |
| C87407_MEL_BA_GR# | SRR40711693 | *S. melanogaster* | BA | GR | M | 98,685,393 | 85.0 | 72.3 | 70.8 |
| C87407_MEL_BR_BC | SRR40711692 | *S. melanogaster* | BR | BC | M | 97,266,526 | 87.2 | 71.5 | 71.2 |
| C87407_MEL_CR_GR | SRR40711691 | *S. melanogaster* | CR | GR | M | 91,038,199 | 73.7 | 70.3 | 56.7 |
| C92298_BOU_CR_BC# | SRR40711690 | *S. bouvreuil* | CR | BC | M | 82,328,798 | 72.9 | 68.1 | 49.8 |
| C92299_MEL_BR_BC | SRR40711689 | *S. melanogaster* | BR | BC | M | 92,402,888 | 87.1 | 70.4 | 66.7 |
| C95860_HYP_BR_OR | SRR40711688 | *S. hypoxantha* | BR | OR | M | 87,113,725 | 86.0 | 69.1 | 63.4 |
| C95860_HYP_CR_GR | SRR40711687 | *S. hypoxantha* | CR | GR | M | 66,878,709 | 74.5 | 61.5 | 38.9 |

| | | | | | | | | | |
|---|---|---|---|---|---|---|---|---|---|
| D131229_SP_BR_BU | SRR40711686 | *Sporophila sp.* | BR | BR | F | 84,242,318 | 76.3 | 65.5 | 51.9 |
| D131229_SP_CR_BO | SRR40711684 | *Sporophila sp.* | CR | BR | F | 78,564,193 | 76.1 | 67.8 | 48.9 |
| D131230_SP_CR_BO | SRR40711683 | *Sporophila sp.* | CR | BR | F | 95,968,298 | 80.1 | 68.0 | 64.6 |
| D147537_BOU_CR_BC | SRR40711682 | *S. bouvreuil* | CR | BC | M | 77,167,674 | 81.4 | 71.7 | 53.1 |
| D165854_SP_CR_BR* | SRR40711681 | *Sporophila sp.* | CR | BR | F | 62,892,907 | 56.5 | 40.0 | 18.6 |
| D165869_PIL_BR_SA | SRR40711680 | *S. pileata* | BR | SA | M | 67,381,141 | 47.1 | 66.7 | 25.9 |
| D165875_MEL_BA_GR | SRR40711679 | *S. melanogaster* | BA | GR | M | 73,634,123 | 74.7 | 69.8 | 45.6 |
| D165875_MEL_BR_BC[#] | SRR40711678 | *S. melanogaster* | BR | BC | M | 73,522,039 | 70.4 | 66.4 | 41.5 |
| D165878_PIL_BA_SA[#] | SRR40711677 | *S. pileata* | BA | SA | M | 82,951,102 | 81.8 | 62.4 | 53.1 |
| D165878_PIL_CR_BC | SRR40711676 | *S. pileata* | CR | BC | M | 87,643,987 | 69.5 | 65.2 | 50.0 |
| D165880_MEL_CR_GR | SRR40711675 | *S. melanogaster* | CR | GR | M | 73,765,130 | 77.0 | 62.8 | 46.0 |
| D165890_PIL_BA_SA | SRR40711673 | *S. pileata* | BA | SA | M | 76,673,769 | 69.3 | 62.8 | 42.2 |
| D165892_PIL_BA_SA | SRR40711672 | *S. pileata* | BA | SA | M | 46,383,248 | 76.5 | 65.9 | 28.4 |
| D179002_MEL_BA_GR | SRR40711671 | *S. melanogaster* | BA | GR | M | 74,958,960 | 79.1 | 64.3 | 47.8 |
| D179002_MEL_BR_BC | SRR40711670 | *S. melanogaster* | BR | BC | M | 94,625,812 | 81.5 | 63.9 | 62.4 |
| D179002_MEL_CR_GR[#] | SRR40711669 | *S. melanogaster* | CR | GR | M | 90,716,292 | 71.3 | 68.0 | 54.0 |
| D179004_MEL_BA_GR | SRR40711668 | *S. melanogaster* | BA | GR | M | 78,757,488 | 81.0 | 72.1 | 53.6 |
| D179004_MEL_BR_BC | SRR40711667 | *S. melanogaster* | BR | BC | M | 72,825,356 | 70.5 | 66.9 | 42.1 |
| D179007_PIL_BA_SA | SRR40711666 | *S. pileata* | BA | SA | M | 92,497,663 | 84.1 | 69.7 | 64.5 |
| D179008_HYP_BR_OR | SRR40711665 | *S. hypoxantha* | BR | OR | M | 90,584,000 | 87.5 | 71.1 | 66.4 |
| D179010_PIL_BA_SA | SRR40711664 | *S. pileata* | BA | SA | M | 91,313,920 | 85.1 | 69.4 | 65.1 |
| D179025_MEL_BR_BC | SRR40711662 | *S. melanogaster* | BR | BC | M | 79,952,810 | 84.6 | 68.3 | 56.2 |
| D179027_PAL_BE_OR[#] | SRR40711661 | *S. palustris* | BR | OR | M | 92,483,004 | 78.2 | 65.7 | 59.1 |
| D179028_SP_BA_BO[#] | SRR40711660 | *Sporophila sp.* | BA | BR | M | 74,940,564 | 75.8 | 71.0 | 47.6 |
| D179028_SP_CR_BO | SRR40711659 | *Sporophila sp.* | CR | BR | M | 79,720,261 | 63.7 | 68.7 | 41.5 |
| D179032_MEL_BR_BC | SRR40711658 | *S. melanogaster* | BR | BC | M | 95,146,092 | 83.6 | 70.5 | 66.6 |
| D179036_PIL_BE_SA | SRR40711657 | *S. pileata* | BR | SA | M | 88,410,903 | 84.5 | 71.5 | 63.6 |
| A09750_HYP_BE_BR | SRR40711655 | *S. hypoxantha* | BR | BR | F | 81,465,146 | 77.1 | 22.0 | |
| A09751_HYP_BA_GR | SRR40711654 | *S. hypoxantha* | BA | GR | M | 87,587,703 | 75.3 | 21.4 | |

| | | | | | | | | |
|---|---|---|---|---|---|---|---|---|
| A09751_HYP_BE_OR | SRR40711653 | *S. hypoxantha* | BR | OR | M | 80,889,695 | 72.6 | 9.6 |
| A09753_SP_BA_BR | SRR40711651 | *Sporophila sp.* | BA | BR | UNK | 79,768,696 | 68.0 | 21.6 |
| A09753_SP_BE_BR | SRR40711650 | *Sporophila sp.* | BR | BR | UNK | 80,225,199 | 72.3 | 23.7 |
| A09753_SP_CR_BR | SRR40711649 | *Sporophila sp.* | CR | BR | UNK | 81,213,566 | 66.2 | 21.2 |
| A09764_CIN_CR_GR | SRR40711648 | *S. cinnamomea* | CR | GR | M | 129,798,344 | 81.0 | 30.3 |
| A09766_CIN_BE_CH | SRR40711647 | *S. cinnamomea* | BR | CH | M | 81,088,687 | 76.9 | 25.7 |
| A09766_CIN_BR_CH | SRR40711646 | *S. cinnamomea* | BR | CH | M | 84,188,659 | 72.7 | 26.7 |
| A09802_HYP_BA_GR | SRR40711645 | *S. hypoxantha* | BA | GR | M | 69,324,220 | 80.3 | 4.0 |
| A09802_HYP_BE_OR | SRR40711644 | *S. hypoxantha* | BR | OR | M | 86,029,170 | 79.4 | 26.0 |

**Table S2. Number of differentially expressed genes in each pairwise patch contrast across four experimental designs.** We tested four designs: primary (~ sex + color + patch), uncontrolled (~ patch), males-only (~ color + patch), and paired (~ individual + patch, restricted to the 11 individuals sampled at more than one patch). Each design involved testing models for each pairwise contrast among crown (CR), breast (BR), and back (BA). The table gives the number of differentially expressed genes for each statistical test.

| Model | Contrast | N DE |
|---|---|---|
| ~ sex + color + patch (primary) | CR vs BR | 70 |
| ~ sex + color + patch (primary) | CR vs BA | 77 |
| ~ sex + color + patch (primary) | BR vs BA | 11 |
| ~ patch | CR vs BR | 57 |
| ~ patch | CR vs BA | 66 |
| ~ patch | BR vs BA | 14 |
| ~ color + patch (males) | CR vs BR | 97 |
| ~ color + patch (males) | CR vs BA | 90 |
| ~ color + patch (males) | BR vs BA | 8 |
| ~ individual + patch (paired) | CR vs BR | 73 |
| ~ individual + patch (paired) | CR vs BA | 78 |
| ~ individual + patch (paired) | BR vs BA | 15 |

**Table S3. Differentially expressed genes across the three pairwise patch contrasts under the primary model.** For each gene we report the contrasts in which it was significant, the adjusted *p*-value for each pairwise comparison, with "TRUE" indicating that the gene met all significance criteria. The final columns indicate whether each gene was recovered under the uncontrolled, males-only, and paired models (Model details in Table S2). *ASIP* (*) was the only candidate gene differentially expressed.

| Gene | N sig | CR vs BR padj | CR vs BA padj | BR vs BA padj | CR vs BR sig | CR vs BA sig | BR vs BA sig | Recovered uncontrolled | Recovered males only | Recovered paired |
|---|---|---|---|---|---|---|---|---|---|---|
| *HAND2* | 3 | 5.59E-67 | 2.79E-04 | 1.24E-28 | TRUE | TRUE | TRUE | TRUE | TRUE | TRUE |
| *HOXA5* | 2 | 1.50E-111 | 7.92E-73 | 6.18E-01 | TRUE | TRUE | – | TRUE | TRUE | TRUE |
| *HOXA7* | 2 | 1.07E-94 | 1.23E-62 | 8.62E-01 | TRUE | TRUE | – | TRUE | TRUE | TRUE |
| *HOXA6* | 2 | 1.64E-87 | 8.21E-66 | 9.08E-01 | TRUE | TRUE | – | TRUE | TRUE | TRUE |
| *HOXA3* | 2 | 6.60E-86 | 2.18E-50 | 6.30E-01 | TRUE | TRUE | – | TRUE | TRUE | TRUE |
| *HOXA4* | 2 | 1.16E-73 | 2.41E-45 | 7.06E-01 | TRUE | TRUE | – | TRUE | TRUE | TRUE |
| *HOXB7* | 2 | 1.30E-45 | 2.33E-35 | 5.77E-01 | TRUE | TRUE | – | TRUE | TRUE | TRUE |
| *HOXB5* | 2 | 1.30E-45 | 4.77E-47 | 6.62E-01 | TRUE | TRUE | – | TRUE | TRUE | TRUE |
| *HOXB6* | 2 | 2.02E-44 | 1.57E-31 | 9.73E-01 | TRUE | TRUE | – | TRUE | TRUE | TRUE |
| *HOXC6* | 2 | 1.97E-32 | 3.89E-24 | 9.22E-01 | TRUE | TRUE | – | TRUE | TRUE | TRUE |
| *HOXA9* | 2 | 5.38E-23 | 5.19E-24 | 6.16E-01 | TRUE | TRUE | – | TRUE | TRUE | TRUE |
| *HOXD4* | 2 | 4.61E-16 | 1.29E-11 | 9.66E-01 | TRUE | TRUE | – | TRUE | TRUE | TRUE |
| *LOC100190237* | 2 | 1.20E-14 | 4.82E-01 | 3.73E-06 | TRUE | – | TRUE | TRUE | TRUE | TRUE |
| *HOXA2* | 2 | 1.42E-11 | 1.40E-05 | 7.72E-01 | TRUE | TRUE | – | TRUE | TRUE | TRUE |
| *ASIP** | 2 | 1.92E-10 | 2.89E-05 | 8.11E-01 | TRUE | TRUE | – | TRUE | TRUE | TRUE |
| *MBP* | 2 | 2.83E-10 | 2.60E-04 | 6.91E-01 | TRUE | TRUE | – | TRUE | TRUE | TRUE |
| *TBX3* | 2 | 4.50E-09 | 8.90E-01 | 6.89E-05 | TRUE | – | TRUE | TRUE | TRUE | TRUE |
| *HPSE2* | 2 | 4.50E-09 | 2.90E-01 | 1.61E-10 | TRUE | – | TRUE | TRUE | TRUE | TRUE |
| *CAPSL* | 2 | 4.42E-07 | 4.31E-04 | 8.94E-01 | TRUE | TRUE | – | TRUE | TRUE | TRUE |
| *g_11493* | 2 | 4.61E-07 | 1.08E-05 | 9.20E-01 | TRUE | TRUE | – | TRUE | TRUE | TRUE |
| *RNLS* | 2 | 3.26E-06 | 2.77E-04 | 9.89E-01 | TRUE | TRUE | – | TRUE | TRUE | – |
| *TMPRSS4* | 2 | 1.12E-05 | 2.14E-06 | 7.23E-01 | TRUE | TRUE | – | TRUE | TRUE | – |
| *NPM2* | 2 | 3.07E-05 | 2.63E-04 | 9.43E-01 | TRUE | TRUE | – | TRUE | TRUE | – |

| | | | | | | | | | | |
|---|---|---|---|---|---|---|---|---|---|---|
| *TPPP3* | 2 | 3.68E-05 | 1.14E-04 | 8.83E-01 | TRUE | TRUE | – | TRUE | TRUE | TRUE |
| *GPR37* | 2 | 5.06E-05 | 3.03E-04 | 9.25E-01 | TRUE | TRUE | – | TRUE | TRUE | – |
| *CORIN* | 2 | 1.73E-04 | 2.24E-04 | 8.46E-01 | TRUE | TRUE | – | TRUE | TRUE | TRUE |
| *NETO2* | 2 | 1.78E-04 | 8.48E-04 | 9.33E-01 | TRUE | TRUE | – | – | TRUE | – |
| *LVRN* | 2 | 2.09E-04 | 2.80E-03 | 9.99E-01 | TRUE | TRUE | – | TRUE | – | TRUE |
| *LOC100225830* | 2 | 4.58E-04 | 4.62E-01 | 2.35E-04 | TRUE | – | TRUE | TRUE | TRUE | – |
| *TYMP* | 2 | 5.15E-04 | 4.76E-03 | 9.96E-01 | TRUE | TRUE | – | – | TRUE | – |
| *LOC101747868* | 2 | 8.34E-04 | 3.22E-04 | 7.83E-01 | TRUE | TRUE | – | TRUE | TRUE | – |
| *SLC5A8* | 2 | 9.33E-04 | 5.22E-08 | 2.18E-01 | TRUE | TRUE | – | TRUE | TRUE | TRUE |
| *SIM1* | 2 | 5.37E-03 | 1.52E-04 | 5.90E-01 | TRUE | TRUE | – | TRUE | TRUE | TRUE |
| *ENPP2* | 2 | 1.28E-01 | 5.17E-09 | 6.91E-04 | – | TRUE | TRUE | TRUE | TRUE | TRUE |
| *ZIC1* | 2 | 2.78E-01 | 1.51E-06 | 6.10E-03 | – | TRUE | TRUE | TRUE | TRUE | TRUE |
| *ALX1* | 2 | 7.92E-01 | 5.01E-07 | 1.01E-07 | – | TRUE | TRUE | TRUE | TRUE | TRUE |
| *SDC3* | 1 | 2.40E-14 | 3.83E-05 | 5.31E-01 | TRUE | – | – | TRUE | TRUE | TRUE |
| *LOC100221249* | 1 | 3.31E-11 | 1.41E-02 | 2.02E-01 | TRUE | – | – | TRUE | TRUE | TRUE |
| *SLC18A2* | 1 | 2.61E-10 | 4.83E-02 | 1.39E-01 | TRUE | – | – | TRUE | TRUE | TRUE |
| *FMO5* | 1 | 2.23E-09 | 1.16E-13 | 3.33E-01 | – | TRUE | – | TRUE | TRUE | TRUE |
| *ENPEP* | 1 | 5.84E-09 | 8.70E-02 | 1.39E-01 | TRUE | – | – | TRUE | TRUE | TRUE |
| *SLC7A2* | 1 | 8.84E-08 | 2.92E-04 | 8.67E-01 | TRUE | – | – | – | TRUE | – |
| *SLC9A9* | 1 | 1.77E-07 | 1.20E-03 | 7.90E-01 | TRUE | – | – | – | TRUE | – |
| *HOXA10* | 1 | 7.65E-07 | 2.41E-02 | 3.56E-01 | TRUE | – | – | – | TRUE | – |
| *P4HA1* | 1 | 9.08E-07 | 1.88E-03 | 8.13E-01 | TRUE | – | – | – | TRUE | TRUE |
| *LOC107053353* | 1 | 2.53E-06 | 1.31E-02 | 6.70E-01 | TRUE | – | – | TRUE | TRUE | TRUE |
| *CPN1* | 1 | 3.16E-06 | 4.89E-03 | 7.82E-01 | TRUE | – | – | TRUE | TRUE | TRUE |
| *IL31RA* | 1 | 3.31E-06 | 2.48E-02 | 6.15E-01 | TRUE | – | – | TRUE | TRUE | – |
| *ADAM19* | 1 | 4.29E-06 | 4.17E-01 | 1.17E-01 | TRUE | – | – | – | TRUE | – |
| *LOC100232284* | 1 | 9.93E-06 | 3.05E-01 | 1.99E-01 | TRUE | – | – | TRUE | TRUE | – |
| *LOC100230413* | 1 | 1.79E-05 | 7.97E-01 | 2.90E-02 | TRUE | – | – | TRUE | TRUE | – |
| *COL21A1* | 1 | 2.03E-05 | 2.39E-10 | 2.24E-01 | – | TRUE | – | TRUE | TRUE | TRUE |

| | | | | | | | | | | |
|---|---|---|---|---|---|---|---|---|---|---|
| *ADAMTS12* | 1 | 4.10E-05 | 2.73E-06 | 6.83E-01 | – | TRUE | – | TRUE | TRUE | TRUE |
| *KANK4* | 1 | 4.17E-05 | 9.43E-02 | 5.35E-01 | TRUE | – | – | – | – | – |
| *ADGRB3* | 1 | 4.36E-05 | 1.87E-01 | 3.75E-01 | TRUE | – | – | – | TRUE | – |
| *CRABP1* | 1 | 4.80E-05 | 1.31E-02 | 8.06E-01 | TRUE | – | – | – | TRUE | TRUE |
| *STC1* | 1 | 4.80E-05 | 5.61E-02 | 6.32E-01 | TRUE | – | – | TRUE | TRUE | – |
| *LUM* | 1 | 4.85E-05 | 1.05E-03 | 9.96E-01 | TRUE | – | – | TRUE | TRUE | TRUE |
| *LOC418114* | 1 | 8.20E-05 | 1.61E-03 | 9.95E-01 | TRUE | – | – | – | TRUE | – |
| *OMD* | 1 | 8.37E-05 | 4.47E-01 | 2.05E-01 | TRUE | – | – | – | – | – |
| *MDK* | 1 | 8.84E-05 | 6.42E-05 | 8.09E-01 | – | TRUE | – | – | TRUE | TRUE |
| *ADAMTS3* | 1 | 9.11E-05 | 9.91E-02 | 5.82E-01 | TRUE | – | – | – | TRUE | – |
| *ADAM12* | 1 | 1.29E-04 | 3.39E-01 | 2.99E-01 | TRUE | – | – | TRUE | TRUE | – |
| *ADORA2B* | 1 | 1.56E-04 | 3.15E-02 | 7.68E-01 | TRUE | – | – | – | – | TRUE |
| *EMX2* | 1 | 1.86E-04 | 3.20E-01 | 3.50E-01 | TRUE | – | – | TRUE | TRUE | TRUE |
| *OTX1* | 1 | 1.94E-04 | 1.38E-01 | 5.75E-01 | TRUE | – | – | TRUE | TRUE | TRUE |
| *EDYM1* | 1 | 1.98E-04 | 1.51E-02 | 8.71E-01 | TRUE | – | – | TRUE | TRUE | TRUE |
| *WDR72* | 1 | 2.64E-04 | 2.77E-01 | 4.15E-01 | TRUE | – | – | TRUE | TRUE | – |
| *ST6GAL2* | 1 | 2.76E-04 | 2.57E-05 | 6.92E-01 | – | TRUE | – | TRUE | TRUE | TRUE |
| *g_13493* | 1 | 3.28E-04 | 6.18E-04 | 8.85E-01 | – | TRUE | – | TRUE | – | TRUE |
| *CPNE9* | 1 | 3.31E-04 | 5.40E-02 | 7.50E-01 | TRUE | – | – | – | – | TRUE |
| *GPC2* | 1 | 3.47E-04 | 5.74E-09 | 1.99E-01 | – | TRUE | – | TRUE | TRUE | TRUE |
| *g_19927* | 1 | 3.59E-04 | 3.39E-01 | 3.85E-01 | TRUE | – | – | – | TRUE | – |
| *LOC100225730* | 1 | 4.49E-04 | 7.31E-01 | 1.43E-01 | TRUE | – | – | – | TRUE | – |
| *MYCN* | 1 | 5.48E-04 | 3.88E-05 | 6.69E-01 | – | TRUE | – | – | TRUE | – |
| *FRRS1L* | 1 | 5.68E-04 | 5.88E-02 | 7.71E-01 | TRUE | – | – | – | TRUE | – |
| *TIMD4* | 1 | 5.95E-04 | 1.49E-01 | 6.30E-01 | TRUE | – | – | – | TRUE | – |
| *RELL2* | 1 | 6.05E-04 | 1.30E-02 | 9.43E-01 | TRUE | – | – | TRUE | TRUE | – |
| *SERPINH1* | 1 | 9.40E-04 | 1.01E-04 | 6.92E-01 | – | TRUE | – | TRUE | TRUE | TRUE |
| *CRHR1* | 1 | 1.30E-03 | 8.81E-02 | 7.71E-01 | TRUE | – | – | TRUE | TRUE | TRUE |
| *DRAXIN* | 1 | 1.78E-03 | 3.64E-05 | 5.85E-01 | – | TRUE | – | TRUE | TRUE | TRUE |

| | | | | | | | | | | |
|---|---|---|---|---|---|---|---|---|---|---|
| *IL1RN* | 1 | 1.91E-03 | 1.36E-04 | 6.56E-01 | – | TRUE | – | – | TRUE | TRUE |
| *SVEP1* | 1 | 2.09E-03 | 5.05E-01 | 2.14E-03 | – | – | TRUE | – | TRUE | – |
| *P4HA2* | 1 | 2.44E-03 | 1.97E-02 | 9.81E-01 | TRUE | – | – | – | TRUE | – |
| *LRRC72* | 1 | 2.79E-03 | 9.38E-06 | 4.34E-01 | – | TRUE | – | TRUE | TRUE | – |
| *PXDNL* | 1 | 3.91E-03 | 4.57E-04 | 7.02E-01 | – | TRUE | – | TRUE | TRUE | TRUE |
| *VIT* | 1 | 4.91E-03 | 6.86E-01 | 3.51E-01 | TRUE | – | – | – | – | TRUE |
| *UGT8* | 1 | 6.65E-03 | 3.29E-04 | 6.28E-01 | – | TRUE | – | – | TRUE | TRUE |
| *LOC100225636* | 1 | 8.96E-03 | 5.72E-04 | 6.40E-01 | – | TRUE | – | TRUE | TRUE | – |
| *g_18725* | 1 | 1.41E-02 | 7.64E-08 | 8.03E-02 | – | TRUE | – | TRUE | TRUE | TRUE |
| *FOS* | 1 | 1.63E-02 | 5.63E-07 | 1.35E-01 | – | TRUE | – | – | TRUE | TRUE |
| *g_16729* | 1 | 1.78E-02 | 5.45E-06 | 2.08E-01 | – | TRUE | – | TRUE | TRUE | TRUE |
| *SLC16A6* | 1 | 2.26E-02 | 6.42E-04 | 5.55E-01 | – | TRUE | – | TRUE | TRUE | – |
| *g_8101* | 1 | 2.55E-02 | 2.49E-04 | 4.48E-01 | – | TRUE | – | TRUE | TRUE | – |
| *LOC100221042* | 1 | 2.79E-02 | 1.42E-03 | 6.04E-01 | – | TRUE | – | – | TRUE | TRUE |
| *SLC22A5* | 1 | 2.86E-02 | 5.42E-04 | 5.10E-01 | – | TRUE | – | – | TRUE | – |
| *FAM162B* | 1 | 4.06E-02 | 1.03E-05 | 1.64E-01 | – | TRUE | – | TRUE | TRUE | – |
| *PTGS2* | 1 | 4.16E-02 | 1.44E-03 | 5.54E-01 | – | TRUE | – | – | TRUE | – |
| *CDO1* | 1 | 5.13E-02 | 1.28E-04 | 2.92E-01 | – | TRUE | – | TRUE | TRUE | TRUE |
| *HSD11B2* | 1 | 5.23E-02 | 1.40E-03 | 5.23E-01 | – | TRUE | – | – | TRUE | – |
| *LOC417013* | 1 | 5.35E-02 | 1.97E-04 | 3.16E-01 | – | TRUE | – | TRUE | TRUE | – |
| *C5H14orf37* | 1 | 6.94E-02 | 1.22E-06 | 5.87E-02 | – | TRUE | – | TRUE | TRUE | TRUE |
| *HOXD8* | 1 | 9.20E-02 | 2.31E-05 | 1.35E-01 | – | TRUE | – | TRUE | TRUE | TRUE |
| *GPC3* | 1 | 9.40E-02 | 5.95E-05 | 1.68E-01 | – | TRUE | – | TRUE | TRUE | TRUE |
| *CTNNA2* | 1 | 9.94E-02 | 5.82E-02 | 1.28E-03 | – | – | TRUE | TRUE | – | TRUE |
| *LOC101751614* | 1 | 1.25E-01 | 4.74E-03 | 5.28E-01 | – | TRUE | – | TRUE | – | TRUE |
| *LOC100217864* | 1 | 1.34E-01 | 3.65E-06 | 4.51E-02 | – | TRUE | – | TRUE | TRUE | TRUE |
| *ADAMTSL2L* | 1 | 1.74E-01 | 4.19E-04 | 2.09E-01 | – | TRUE | – | TRUE | TRUE | – |
| *GRIN3A* | 1 | 1.83E-01 | 4.83E-02 | 3.10E-03 | – | – | TRUE | – | TRUE | – |
| *FHL1* | 1 | 1.83E-01 | 2.01E-03 | 3.43E-01 | – | TRUE | – | TRUE | TRUE | TRUE |

| | | | | | | | | | | |
|---|---|---|---|---|---|---|---|---|---|---|
| *NDP* | 1 | 1.85E-01 | 4.58E-05 | 9.62E-02 | – | TRUE | – | TRUE | TRUE | TRUE |
| *LOC105758853* | 1 | 2.09E-01 | 1.61E-03 | 2.93E-01 | – | TRUE | – | – | TRUE | – |
| *nbis-gene-183* | 1 | 2.11E-01 | 5.87E-03 | 4.43E-01 | – | TRUE | – | – | TRUE | – |
| *RBM47* | 1 | 2.22E-01 | 9.29E-06 | 3.02E-02 | – | TRUE | – | – | TRUE | – |
| *SLC5A11* | 1 | 2.67E-01 | 6.01E-04 | 1.68E-01 | – | TRUE | – | TRUE | TRUE | – |
| *nbis-gene-1757* | 1 | 3.12E-01 | 2.70E-04 | 1.14E-01 | – | TRUE | – | – | TRUE | – |
| *OTUD1* | 1 | 3.89E-01 | 2.37E-04 | 8.04E-02 | – | TRUE | – | – | TRUE | TRUE |
| *SLC26A9* | 1 | 4.04E-01 | 2.71E-03 | 2.05E-01 | – | TRUE | – | – | TRUE | – |
| *SERPINC1* | 1 | 5.41E-01 | 1.41E-03 | 1.14E-01 | – | TRUE | – | – | – | – |
| *NR4A1* | 1 | 5.97E-01 | 6.71E-04 | 6.84E-02 | – | TRUE | – | – | TRUE | TRUE |
| *LOC107049603* | 1 | 8.49E-01 | 2.50E-03 | 5.87E-02 | – | TRUE | – | – | – | TRUE |

**Table S4. Number of differentially expressed genes in each pairwise color contrast across seven experimental designs**. We tested seven models: the primary patch-controlled design (~ patch + color, males), the uncontrolled design (~ color, males), the full design including all libraries (~ sex + patch + color), and within-patch analyses performed separately for crown, breast, and back (~ color, males), as well as for crown including brown individuals while controlling for sex (~ sex + color). Other details as in Table S2. Color codes are: BC (black), SA (salmon), OR (orange), GR (gray) and BR (brown).

| Model | N libraries | Contrast | N DE |
|---|---|---|---|
| ~ patch + color (males) [primary] | 34 | BC vs SA | 0 |
| ~ patch + color (males) [primary] | 34 | BC vs OR | 3 |
| ~ patch + color (males) [primary] | 34 | BC vs GR | 0 |
| ~ patch + color (males) [primary] | 34 | SA vs OR | 1 |
| ~ patch + color (males) [primary] | 34 | SA vs GR | 16 |
| ~ patch + color (males) [primary] | 34 | OR vs GR | 2 |
| ~ color (males) | 34 | BC vs SA | 3 |
| ~ color (males) | 34 | BC vs OR | 5 |
| ~ color (males) | 34 | BC vs GR | 3 |
| ~ color (males) | 34 | SA vs OR | 4 |
| ~ color (males) | 34 | SA vs GR | 13 |
| ~ color (males) | 34 | OR vs GR | 5 |
| ~ sex + patch + color (all) | 43 | BC vs SA | 1 |
| ~ sex + patch + color (all) | 43 | BC vs OR | 5 |
| ~ sex + patch + color (all) | 43 | BC vs GR | 0 |
| ~ sex + patch + color (all) | 43 | BC vs BR | 0 |
| ~ sex + patch + color (all) | 43 | SA vs OR | 0 |
| ~ sex + patch + color (all) | 43 | SA vs GR | 8 |
| ~ sex + patch + color (all) | 43 | SA vs BR | 0 |
| ~ sex + patch + color (all) | 43 | OR vs GR | 6 |
| ~ sex + patch + color (all) | 43 | OR vs BR | 0 |
| ~ sex + patch + color (all) | 43 | GR vs BR | 0 |
| within crown (males) | 9 | BC vs GR | 0 |
| within breast (males) | 15 | BC vs SA | 1 |
| within breast (males) | 15 | BC vs OR | 2 |
| within breast (males) | 15 | SA vs OR | 3 |
| within back (males) | 10 | SA vs GR | 12 |
| within crown (all libs, ~ sex + color) | 13 | BC vs GR | 2 |
| within crown (all libs, ~ sex + color) | 13 | BC vs BR | 2 |
| within crown (all libs, ~ sex + color) | 13 | GR vs BR | 7 |

**Table S5. Differentially expressed genes in the color contrasts.** The *primary* column indicates genes identified as differentially expressed in the primary model (~ patch + color, males). For the uncontrolled model and the model including all libraries, entries indicate whether a gene was differentially expressed in at least one comparison. For the within-patch models, entries specifying a color contrast indicate the pairwise comparison in which differential expression was detected. Model details in Table S4 and other details as in Table S2. Candidate genes are marked with an asterisk (*). *DCT* marked with (#) was recovered as an outlier candidate gene in Campagna et al. (2017) *Sci Adv*.

| Gene | Primary | Primary contrasts | Uncontrolled | All libraries | Crown all samples. | Within breast (males) | Within back (males) | Within crown (males) |
|---|---|---|---|---|---|---|---|---|
| *ASIP** | – | NA | TRUE | – | – | – | SA vs GR | – |
| *DCT**# | – | NA | TRUE | – | – | – | SA vs GR | – |
| *FKBP5* | TRUE | SA vs GR | – | – | – | – | SA vs GR | – |
| *LOC101234089* | TRUE | BC vs OR; SA vs OR | TRUE | – | – | BC vs OR; SA vs OR | – | – |
| *MAN2C1* | TRUE | SA vs GR | TRUE | TRUE | – | – | SA vs GR | – |
| *ZBTB16* | TRUE | SA vs GR | – | TRUE | – | – | SA vs GR | – |
| *AKAP9* | – | NA | – | TRUE | – | BC vs OR | – | – |
| *ATF3* | – | NA | – | – | – | – | SA vs GR | – |
| *ATP10D* | – | NA | – | – | – | – | SA vs GR | – |
| *CEP104* | – | NA | – | – | – | SA vs OR | – | – |
| *CEP290* | TRUE | BC vs OR | TRUE | TRUE | – | – | – | – |
| *CHMP2A* | – | NA | – | – | – | – | SA vs GR | – |
| *CLVS2* | TRUE | SA vs GR | – | TRUE | – | – | – | – |
| *DNAJA1* | TRUE | SA vs GR | – | TRUE | – | – | – | – |
| *EFCAB5* | – | NA | – | – | – | – | SA vs GR | – |
| *FABP7* | – | NA | – | – | – | – | SA vs GR | – |
| *FRRS1L* | TRUE | SA vs GR | – | TRUE | – | – | – | – |
| *IRF4* | TRUE | SA vs GR | TRUE | – | – | – | – | – |
| *LOC100217745* | TRUE | SA vs GR | – | – | – | – | – | – |
| *LOC100219651* | – | NA | – | – | – | – | SA vs GR | – |
| *LOC100220369* | – | NA | – | – | – | BC vs SA | – | – |

| | | | | | | | | |
|---|---|---|---|---|---|---|---|---|
| *LOC100223651* | – | NA | – | – | – | – | SA vs GR | – |
| *LOC100225021* | TRUE | SA vs GR | – | – | – | – | – | – |
| *LOC107049018* | – | NA | – | – | – | SA vs OR | – | – |
| *LOC107049603* | TRUE | OR vs GR | – | TRUE | – | – | – | – |
| *LOC415324* | TRUE | SA vs GR | TRUE | TRUE | – | – | – | – |
| *P4HA2* | TRUE | SA vs GR | – | – | – | – | – | – |
| *PHACTR3* | TRUE | BC vs OR | TRUE | TRUE | – | – | – | – |
| *REG4* | TRUE | SA vs GR | TRUE | TRUE | – | – | – | – |
| *SV2B* | TRUE | SA vs GR | TRUE | TRUE | – | – | – | – |
| *TYMP* | TRUE | SA vs GR | – | – | – | – | – | – |
| *g_20731* | TRUE | OR vs GR | TRUE | TRUE | – | – | – | – |
| *g_2345* | TRUE | SA vs GR | – | – | – | – | – | – |
| *id-TCRA.gene* | TRUE | SA vs GR | – | – | – | – | – | – |
| *GLDN* | – | NA | – | – | BC vs GR | – | – | – |
| *GAL3ST1* | – | NA | – | – | BC vs GR | – | – | – |
| *LOC101747901* | – | NA | – | – | BC vs BR, GR vs BR | – | – | – |
| *LOC100218707* | – | NA | – | – | BC vs BR, GR vs BR | – | – | – |
| *g_20088* | – | NA | – | – | GR vs BR | – | – | – |
| *LOC100857297* | – | NA | – | – | GR vs BR | – | – | – |
| *AADACL4L5* | – | NA | – | – | GR vs BR | – | – | – |
| *g_2118* | – | NA | – | – | GR vs BR | – | – | – |
| *KCNE4* | – | NA | – | – | GR vs BR | – | – | – |

**Table S6. Number of sex-biased differentially expressed genes.** We tested three experimental designs: primary (~ sex), patch-adjusted (~ patch + sex), and patch- and color-adjusted (~ patch + color + sex). All designs included the 43 samples with male as the reference level. The table shows the number of differentially expressed genes (N DE), split into Z-linked and autosomal genes, with autosomal genes further split by the direction of bias in expression (higher in males or females). The final column shows, for each model, the number of genes that overlap with the 68 genes identified as significant under the primary model. Chromosome classes as in Table S7.

| Model | N DE | N Z linked | N autosomal | Autosomal higher males | Autosomal higher females | Recovered from primary |
|---|---|---|---|---|---|---|
| ~ sex (primary) | 68 | 45 | 23 | 10 | 13 | 68 |
| ~ patch + sex | 65 | 40 | 25 | 12 | 13 | 55 |
| ~ patch + color + sex | 34 | 30 | 4 | 3 | 1 | 23 |

**Table S7. Sex-biased differentially expressed genes.** For each of the 68 genes significant in the primary sex model (~ sex, male reference), the table gives the gene name and the significance under the different model designs (patch: ~patch+ sex, color: ~patch+color+ sex). The table also shows if the gene is autosomal or Z-linked. Other details as in Table S2.

| Gene | Contig | Padj sex | Sig sex | Padj patch | Sig patch | Padj pcolor | Sig pcolor | N models | Chr-class |
|---|---|---|---|---|---|---|---|---|---|
| *BCO2* | 30 | 5.95E-104 | TRUE | 3.99E-94 | TRUE | 4.59E-17 | TRUE | 3 | autosomal |
| *LOC100221697* | 7 | 5.10E-37 | TRUE | 1.05E-33 | TRUE | 3.53E-08 | – | 2 | Z-linked |
| *ATG10* | 35 | 2.02E-31 | TRUE | 7.60E-29 | TRUE | 6.01E-08 | TRUE | 3 | Z-linked |
| *INIP* | 7 | 4.07E-31 | TRUE | 1.75E-27 | – | 6.89E-06 | – | 1 | Z-linked |
| *GNAO1* | 7 | 2.93E-28 | TRUE | 1.17E-26 | TRUE | 4.30E-06 | TRUE | 3 | Z-linked |
| *PTBP3* | 7 | 7.56E-24 | TRUE | 5.67E-23 | TRUE | 3.40E-06 | TRUE | 3 | Z-linked |
| *LOC100217858* | 7 | 7.46E-23 | TRUE | 6.37E-21 | TRUE | 1.44E-05 | TRUE | 3 | Z-linked |
| *ATP5A1* | 41 | 1.75E-22 | TRUE | 1.21E-21 | TRUE | 5.40E-05 | TRUE | 3 | Z-linked |
| *CZH9orf64* | 7 | 6.55E-22 | TRUE | 2.28E-19 | TRUE | 2.37E-03 | – | 2 | Z-linked |
| *UBAP2* | 7 | 3.16E-20 | TRUE | 2.24E-20 | TRUE | 3.01E-05 | TRUE | 3 | Z-linked |
| *FAM172A* | 28 | 1.86E-17 | TRUE | 1.72E-20 | TRUE | 7.41E-05 | TRUE | 3 | Z-linked |
| *FANCG* | 7 | 2.31E-16 | TRUE | 2.94E-14 | TRUE | 4.89E-02 | – | 2 | Z-linked |
| *RPS23* | 35 | 6.00E-16 | TRUE | 6.37E-14 | – | 2.38E-02 | – | 1 | Z-linked |
| *IKBKAP* | 7 | 9.12E-16 | TRUE | 1.30E-15 | TRUE | 2.67E-06 | TRUE | 3 | Z-linked |
| *SNCAIP* | 7 | 3.24E-15 | TRUE | 1.05E-14 | TRUE | 1.26E-03 | TRUE | 3 | Z-linked |
| *TPGS2* | 7 | 7.18E-15 | TRUE | 4.59E-14 | TRUE | 2.02E-02 | – | 2 | Z-linked |
| *ARSB* | 7 | 9.19E-15 | TRUE | 6.22E-13 | TRUE | 4.05E-02 | – | 2 | Z-linked |
| *RNF20* | 43 | 2.94E-14 | TRUE | 6.49E-13 | TRUE | 1.14E-03 | TRUE | 3 | Z-linked |
| *g_18915* | 7 | 1.11E-13 | TRUE | 1.22E-11 | TRUE | 2.37E-03 | TRUE | 3 | Z-linked |
| *ARHGEF39* | 7 | 2.80E-13 | TRUE | 3.79E-11 | – | 4.23E-03 | TRUE | 2 | Z-linked |
| *CCDC171* | 7 | 2.64E-12 | TRUE | 2.25E-12 | – | 5.81E-05 | TRUE | 2 | Z-linked |
| *POC5* | 7 | 3.85E-12 | TRUE | 6.55E-11 | – | 6.24E-02 | – | 1 | Z-linked |
| *HIATL1* | 7 | 4.65E-12 | TRUE | 3.09E-14 | TRUE | 1.61E-02 | – | 2 | Z-linked |
| *CA9* | 7 | 1.28E-11 | TRUE | 4.66E-12 | TRUE | 1.01E-02 | TRUE | 3 | Z-linked |

| | | | | | | | | | |
|---|---|---|---|---|---|---|---|---|---|
| *FRMD3* | 7 | 1.76E-11 | TRUE | 2.98E-11 | TRUE | 4.97E-02 | – | 2 | Z-linked |
| *DNAJA1* | 7 | 1.96E-10 | TRUE | 1.57E-10 | TRUE | 8.30E-02 | – | 2 | Z-linked |
| *CZH18orf25* | 41 | 3.13E-10 | TRUE | 1.04E-10 | TRUE | 1.65E-02 | TRUE | 3 | Z-linked |
| *PDE8A* | 7 | 1.02E-09 | TRUE | 6.00E-08 | TRUE | 4.71E-02 | TRUE | 3 | Z-linked |
| *STARD4* | 28 | 1.27E-09 | TRUE | 4.54E-10 | TRUE | 8.78E-02 | – | 2 | Z-linked |
| *KIAA0020* | 7 | 2.14E-09 | TRUE | 2.53E-09 | TRUE | 1.39E-02 | TRUE | 3 | Z-linked |
| *TRMT10B* | 35 | 2.93E-09 | TRUE | 1.89E-09 | TRUE | 2.98E-01 | – | 2 | Z-linked |
| *GALT* | 7 | 4.59E-09 | TRUE | 5.90E-08 | TRUE | 1.12E-01 | – | 2 | Z-linked |
| *GRHPR* | 35 | 4.81E-09 | TRUE | 1.94E-08 | TRUE | 4.52E-01 | – | 2 | Z-linked |
| *FOXE1* | 7 | 6.92E-09 | TRUE | 8.72E-10 | TRUE | 3.42E-02 | TRUE | 3 | Z-linked |
| *ANXA1* | 28 | 3.28E-08 | TRUE | 6.69E-10 | TRUE | 1.52E-02 | TRUE | 3 | Z-linked |
| *MEX3B* | 41 | 3.87E-07 | TRUE | 2.87E-06 | TRUE | 3.15E-02 | TRUE | 3 | Z-linked |
| *TMEM171* | 7 | 5.55E-07 | TRUE | 1.17E-05 | TRUE | 5.94E-01 | – | 2 | Z-linked |
| *MLANA* | 7 | 3.28E-06 | TRUE | 1.22E-05 | TRUE | 7.29E-01 | – | 2 | Z-linked |
| *GADD45G* | 7 | 5.28E-06 | TRUE | 2.72E-07 | TRUE | 3.30E-01 | – | 2 | Z-linked |
| *MAP1B* | 7 | 1.15E-05 | TRUE | 1.49E-04 | – | 9.14E-02 | – | 1 | Z-linked |
| *g_14776* | 41 | 1.19E-05 | TRUE | 1.46E-05 | TRUE | 2.36E-01 | – | 2 | Z-linked |
| *DDX58* | 7 | 9.63E-05 | TRUE | 4.82E-04 | – | 5.25E-01 | – | 1 | Z-linked |
| *g_2118* | 10 | 1.89E-04 | TRUE | 3.18E-03 | TRUE | 8.81E-01 | – | 2 | autosomal |
| *g_3503* | 120 | 1.95E-04 | TRUE | 2.65E-03 | TRUE | 4.61E-01 | – | 2 | autosomal |
| *g_2860* | 11 | 2.35E-04 | TRUE | 2.14E-06 | TRUE | 2.98E-01 | – | 2 | autosomal |
| *TROJANZ* | 7 | 2.93E-04 | TRUE | 7.86E-04 | – | 4.92E-01 | – | 1 | Z-linked |
| *LOC100218614* | 1 | 3.28E-04 | TRUE | 1.96E-03 | TRUE | 6.08E-01 | – | 2 | autosomal |
| *CNTN4* | 12 | 3.65E-04 | TRUE | 1.04E-02 | TRUE | 6.64E-01 | – | 2 | autosomal |
| *ACTN1* | 1 | 4.17E-04 | TRUE | 1.31E-04 | TRUE | 6.51E-02 | – | 2 | autosomal |
| *ARID3C* | 7 | 4.63E-04 | TRUE | 1.08E-04 | TRUE | 9.81E-02 | – | 2 | Z-linked |
| *KIAA1045* | 7 | 6.66E-04 | TRUE | 2.32E-05 | TRUE | 4.43E-01 | – | 2 | Z-linked |
| *LOC100226706* | 14 | 8.18E-04 | TRUE | 1.86E-03 | TRUE | 7.96E-01 | – | 2 | autosomal |
| *LOC100231107* | 9 | 9.97E-04 | TRUE | 6.64E-04 | TRUE | 5.87E-01 | – | 2 | autosomal |

| | | | | | | | | | |
|---|---|---|---|---|---|---|---|---|---|
| *APOLD1* | 5 | 1.29E-03 | TRUE | 1.23E-03 | TRUE | 6.75E-01 | – | 2 | autosomal |
| *LOC107055264* | 40 | 1.47E-03 | TRUE | 1.01E-03 | TRUE | 1.01E-01 | – | 2 | autosomal |
| *TEX12* | 30 | 1.91E-03 | TRUE | 9.03E-04 | TRUE | 6.23E-01 | – | 2 | autosomal |
| *RAB36* | 22 | 2.78E-03 | TRUE | 3.24E-03 | TRUE | 4.95E-01 | – | 2 | autosomal |
| *LOC417013* | 22 | 3.41E-03 | TRUE | 1.37E-03 | – | 9.09E-01 | – | 1 | autosomal |
| *g_18725* | 7 | 3.74E-03 | TRUE | 5.97E-08 | TRUE | 3.86E-03 | TRUE | 3 | Z-linked |
| *FOS* | 4 | 4.60E-03 | TRUE | 2.71E-05 | TRUE | 1.68E-02 | TRUE | 3 | autosomal |
| *LOC100226251* | 2 | 6.89E-03 | TRUE | 2.40E-03 | TRUE | 5.94E-01 | – | 2 | autosomal |
| *KIAA0319* | 2 | 1.00E-02 | TRUE | 1.53E-03 | TRUE | 2.14E-01 | – | 2 | autosomal |
| *NDUFA4L2* | 197 | 1.08E-02 | TRUE | 4.04E-02 | – | 8.52E-01 | – | 1 | autosomal |
| *RASL10A* | 32 | 1.20E-02 | TRUE | 1.80E-02 | TRUE | 9.82E-01 | – | 2 | autosomal |
| *g_10068* | 27 | 1.29E-02 | TRUE | 1.26E-02 | TRUE | 3.78E-01 | – | 2 | autosomal |
| *LIPE* | 83 | 2.58E-02 | TRUE | 4.24E-02 | – | 7.25E-01 | – | 1 | autosomal |
| *HAUS3L* | 12 | 3.67E-02 | TRUE | 6.01E-02 | – | 8.03E-01 | – | 1 | autosomal |
| *KRT20* | 40 | 4.73E-02 | TRUE | 6.24E-02 | – | 9.40E-01 | – | 1 | autosomal |

**Table S8. Feather carotenoid content by individual.** Concentrations are expressed as µg g$^{-1}$ feather for cis-lutein, lutein, zeaxanthin, other carotenoids, and the total content. Feather samples are derived from adult individuals that are different from those in the gene expression study. Values of 0 indicate concentrations below the detection threshold (2 µg g$^{-1}$). Patch abbreviations: CR (Crown), BR (Breast) and TH (Throat).

| Sample | Species | Sex | Patch | Cis-lutein (µg g$^{-1}$) | Lutein (µg g$^{-1}$) | Zeaxanthin (µg g$^{-1}$) | Other (µg g$^{-1}$) | Total (µg g$^{-1}$) |
|---|---|---|---|---|---|---|---|---|
| BOU_17 | *S. bouvreuil* | M | BR | 0 | 0 | 0 | 0 | 0 |
| BOU_17 | *S. bouvreuil* | M | CR | 0 | 0 | 0 | 0 | 0 |
| BOU_18 | *S. bouvreuil* | F | BR | 2.4 | 3.3 | 2.6 | 0.0 | 8.3 |
| BOU_18 | *S. bouvreuil* | F | CR | 4.9 | 13.1 | 5.3 | 7.9 | 31.1 |
| HX9033 | *S. hypoxantha* | M | BR | 0 | 0 | 0 | 0 | 0 |
| HX9033 | *S. hypoxantha* | M | CR | 0 | 0 | 0 | 0 | 0 |
| HX9051 | *S. hypoxantha* | F | BR | 2.3 | 3.2 | 2.5 | 0.0 | 8.0 |
| HX9051 | *S. hypoxantha* | F | CR | 5.7 | 12.4 | 6.9 | 9.7 | 34.6 |
| HX9066 | *S. hypoxantha* | F | BR | 3.6 | 5.0 | 4.0 | 3.6 | 16.2 |
| HX9066 | *S. hypoxantha* | F | CR | 8.6 | 13.3 | 9.6 | 7.8 | 39.3 |
| HX9078 | *S. hypoxantha* | M | BR | 0 | 0 | 0 | 0 | 0 |
| HX9078 | *S. hypoxantha* | M | CR | 4.7 | 5.7 | 5.1 | 0.0 | 15.5 |
| HYP_30 | *S. hypochroma* | F | BR | 0 | 2.6 | 2.3 | 0 | 5.0 |
| HYP_30 | *S. hypochroma* | F | CR | 5.9 | 15.8 | 6.6 | 9.4 | 37.8 |
| IB9043 | *S. iberaensis* | M | BR | 0 | 0 | 0 | 0 | 0 |
| IB9043 | *S. iberaensis* | M | CR | 0 | 0 | 0.0 | 0 | 0 |
| IB9046 | *S. iberaensis* | M | BR | 0 | 0 | 0 | 0 | 0 |
| IB9046 | *S. iberaensis* | M | CR | 0 | 3.1 | 0.0 | 0 | 3.1 |
| IB9501 | *S. iberaensis* | F | BR | 2.3 | 3.0 | 2.8 | 0 | 8.1 |
| IB9501 | *S. iberaensis* | F | CR | 8.8 | 14.4 | 11.0 | 8.0 | 42.2 |
| IB9586 | *S. iberaensis* | F | BR | 2.8 | 4.0 | 3.5 | 0 | 10.2 |
| IB9586 | *S. iberaensis* | F | CR | 5.3 | 10.0 | 7.0 | 9.1 | 31.4 |
| MEL_25 | *S. melanogaster* | M | CR | 0 | 0 | 0 | 0 | 0 |

| | | | | | | | | |
|---|---|---|---|---|---|---|---|---|
| MEL_3 | *S. melanogaster* | M | BR | 0 | 0 | 0 | 0 | 0 |
| PAL_6 | *S. palustris* | M | BR | 0 | 5.4 | 0 | 0 | 5.4 |
| PAL_6 | *S. palustris* | M | CR | 0 | 6.8 | 0 | 0 | 6.8 |
| PAL_6 | *S. palustris* | M | TH | 0 | 5.3 | 0 | 0 | 5.3 |
| RUF_1 | *S. ruficollis* | F | BR | 3.5 | 11.8 | 3.8 | 4.9 | 24.1 |
| RUF_1 | *S. ruficollis* | F | CR | 7.0 | 15.1 | 7.9 | 11.5 | 41.4 |

**Table S9. Predicted consequences and species allele frequencies of outlier variants within candidate pigmentation genes.** For each $F_{ST}$/GWAS outlier variant falling within a candidate pigmentation gene, we report coordinates on the *S. hypoxantha* reference genome, the reference (ref) and alternate (alt) alleles, the variant type (snp or indel), and the consequence predicted with bcftools csq under the improved annotation. For the amino-acid change for coding variants (aa change), those marked with an asterisk alter the protein sequence. Protein-changing variants are reported against the isoform with the highest mean expression among those carrying them (transcript), with that isoform's share of gene expression given as transcript % (see Methods). Variants comprise missense (8), synonymous (7), splice region (3), 5′ UTR (18), and 3′ UTR (33). The frequency columns show the alternate allele frequency in each species (bou, *S. bouvreuil*, n = 15; cin, *S. cinnamomea*, 15; hypoch, *S. hypochroma*, 17; hypox, *S. hypoxantha*, 22; ibe, *S. iberaensis*, 19; mel, *S. melanogaster*, 9; nig, *S. nigrorufa*, 4; pal, *S. palustris*, 7; pil, *S. pileata*, 6; ruf, *S. ruficollis*, 13); NA indicates no reads at that position in that species.

| | | | | | | | Alternate allele frequency | | | | | | | | | |
|---|---|---|---|---|---|---|---|---|---|---|---|---|---|---|---|---|
| **Gene** | **Contig: position** | **Ref/alt** | **Variant type** | **Consequence** | **aa change** | **Transcript %** | **bou** | **cin** | **hypoch** | **hypox** | **ibe** | **mel** | **nig** | **pal** | **pil** | **ruf** |
| *ASIP* | 21:13594449 | T/C | snp | 5′ UTR | NA | 3.3 | 0.93 | 0.31 | 0.01 | 0 | 0.39 | 0 | 0 | 0 | 0.75 | 0 |
| *ASIP* | 21:13594471 | T/C | snp | 5′ UTR | NA | 3.3 | 1 | 0.80 | 0.05 | 0 | 0.63 | 0 | 0 | 0 | 1 | 0 |
| *ASIP* | 21:13604095 | C/T | snp | 5′ UTR | NA | 2.2 | 0 | 0 | 0.03 | 0.13 | 0.05 | 1 | 0 | 0.09 | 0 | 0.07 |
| *ASIP* | 21:13604083 | G/T | snp | 5′ UTR | NA | 2.2 | 0 | 0 | 0.12 | 0.16 | 0.07 | 1 | 0 | 0.09 | 0.04 | 0.07 |
| *ASIP* | 21:13594585 | G/A | snp | 5′ UTR | NA | 3.3 | 0 | 0 | 0.17 | 0.25 | 0.25 | 1 | 1 | 0.25 | 0 | 0.10 |
| *ASIP* | 21:13629967 | G/T | snp | 3′ UTR | NA | 49.9 | 0.42 | 0.22 | 0.25 | 0.19 | 0.27 | 0.91 | 0.40 | 0.10 | 0.74 | 0.25 |
| *ASIP* | 21:13594565 | A/C | snp | 5′ UTR | NA | 3.3 | 0.01 | 0.02 | 0.45 | 0.47 | 0.32 | 1 | 1 | 0.54 | 0.03 | 0.26 |
| *ASIP* | 21:13603978 | T/C | snp | 5′ UTR | NA | 2.2 | 0.96 | 0.83 | 0.11 | 0.04 | 0.34 | 0 | 0.05 | 0.15 | 1 | 0.28 |
| *ASIP* | 21:13604128 | C/A | snp | 5′ UTR | NA | 2.2 | 1 | 0.74 | 0.06 | 0.13 | 0.38 | 0 | 0.08 | 0.15 | 1 | 0.32 |
| *ASIP* | 21:13604173 | C/G | snp | 5′ UTR | NA | 2.2 | 0.95 | 0.52 | 0.19 | 0.13 | 0.41 | 0 | 0 | 0.17 | 0.94 | 0.34 |
| *ASIP* | 21:13615196 | C/T | snp | 5′ UTR | NA | 5.7 | 0.98 | 0.41 | 0.41 | 0.42 | 0.41 | 0 | NA | 0.70 | 1 | 0.42 |
| *ASIP* | 21:13615144 | A/G | snp | 5′ UTR | NA | 5.7 | 0.96 | 0.42 | 0.43 | 0.33 | 0.39 | 0 | NA | 0.67 | 1 | 0.43 |
| *ASIP* | 21:13615066 | T/G | snp | 5′ UTR | NA | 5.7 | 0.97 | 0.45 | 0.46 | 0.39 | 0.42 | 0 | NA | 0.75 | 1 | 0.43 |
| *ASIP* | 21:13615183 | T/C | snp | 5′ UTR | NA | 5.7 | 0.84 | 0.44 | 0.41 | 0.40 | 0.39 | 0 | NA | 0.70 | 1 | 0.44 |
| *ASIP* | 21:13615123 | C/T | snp | 5′ UTR | NA | 5.7 | 0.97 | 0.44 | 0.44 | 0.42 | 0.45 | 0 | NA | 0.67 | 1 | 0.46 |
| *GPT2* | 14:13959589 | G/T | snp | 3′ UTR | NA | 96.2 | 0.75 | 0 | 0.04 | 0.03 | 0.78 | 0 | 1 | 0 | 0.97 | 0 |
| *GPT2* | 14:13959544 | A/AT | indel | 3′ UTR | NA | 96.2 | 0.80 | 0 | 0.05 | 0.03 | 0.74 | 0 | 0.75 | 0 | 1 | 0 |

| | | | | | | | | | | | | | | | |
|---|---|---|---|---|---|---|---|---|---|---|---|---|---|---|---|
| *GPT2* | 14:13959441 | A/G | snp | 3′ UTR | NA | 96.2 | 0.85 | 0 | 0.06 | 0.01 | 0.72 | 0 | 0.89 | 0 | 1 | 0 |
| *GPT2* | 14:13959224 | G/GTTTGT | indel | 3′ UTR | NA | 96.2 | 0.80 | 0.02 | 0.07 | 0 | 0.74 | 0 | 0.90 | 0 | 1 | 0 |
| *GPT2* | 14:13958971 | T/A | snp | 3′ UTR | NA | 96.2 | 0.85 | 0 | 0.02 | 0.02 | 0.69 | 0 | 1 | 0 | 1 | 0 |
| *GPT2* | 14:13958946 | A/C | snp | 3′ UTR | NA | 96.2 | 0.85 | 0 | 0.03 | 0.02 | 0.69 | 0 | 1 | 0 | 1 | 0 |
| *GPT2* | 14:13964245 | G/T | snp | missense | 403L>403I* | 96.2 | 0.77 | 0 | 0.05 | 0 | 0.74 | 0 | 1 | 0 | 1 | 0 |
| *GPT2* | 14:13959690 | T/C | snp | 3′ UTR | NA | 96.2 | 0.76 | 0 | 0.08 | 0.03 | 0.76 | 0 | 1 | 0 | 1 | 0 |
| *GPT2* | 14:13958963 | T/C | snp | 3′ UTR | NA | 96.2 | 0.83 | 0.01 | 0.02 | 0.02 | 0.68 | 0.02 | 1 | 0 | 1 | 0 |
| *GPT2* | 14:13959257 | A/T | snp | 3′ UTR | NA | 96.2 | 0.79 | 0 | 0.04 | 0 | 0.75 | 0.02 | 1 | 0 | 1 | 0 |
| *GPT2* | 14:13959067 | A/T | snp | 3′ UTR | NA | 96.2 | 0.79 | 0 | 0.04 | 0.03 | 0.67 | 0 | 1 | 0 | 1 | 0.01 |
| *GPT2* | 14:13959936 | T/C | snp | 3′ UTR | NA | 96.2 | 0.84 | 0.02 | 0.10 | 0.03 | 0.70 | 0 | 1 | 0 | 1 | 0.01 |
| *GPT2* | 14:13959932 | T/C | snp | 3′ UTR | NA | 96.2 | 0.84 | 0.02 | 0.10 | 0.02 | 0.67 | 0 | 1 | 0 | 1 | 0.01 |
| *GPT2* | 14:13960644 | C/G | snp | 3′ UTR | NA | 96.2 | 0.81 | 0 | 0.05 | 0.02 | 0.73 | 0 | 1 | 0 | 1 | 0.04 |
| *GPT2* | 14:13959707 | G/A | snp | 3′ UTR | NA | 96.2 | 0.80 | 0.31 | 0.31 | 0.13 | 0.88 | 0.12 | 1 | 0.20 | 1 | 0.08 |
| *GPT2* | 14:13959415 | G/A | snp | 3′ UTR | NA | 96.2 | 0.79 | 0.05 | 0.01 | 0.04 | 0.72 | 0.06 | 1 | 0 | 1 | 0.08 |
| *GPT2* | 14:13965772 | C/T | snp | synonymous | 308K | 2.5 | 0.87 | 0.07 | 0.22 | 0.16 | 0.85 | 0.19 | 1 | 0.13 | 1 | 0.09 |
| *GPT2* | 14:13961077 | A/G | snp | 3′ UTR | NA | 96.2 | 0.21 | 0.80 | 0.78 | 0.92 | 0.32 | 0.94 | 0.22 | 0.91 | 0 | 0.81 |
| *HERC2* | 10:6204110 | G/A | snp | synonymous | 1574L | 44.9 | 0.18 | 0.30 | 0.30 | 0.11 | 0.20 | 0.34 | 0 | 0 | 0.10 | 0.88 |
| *HERC2* | 10:6218697 | G/A | snp | 3′ UTR | NA | 4.3 | 0 | 0.05 | 0.17 | 0.02 | 0.28 | 0 | 0 | 0 | 0 | 0.90 |
| *HERC2* | 10:6206235 | A/G | snp | missense | 1727Y>1727C* | 4.3 | 0.89 | 0.67 | 0.33 | 0.06 | 0.67 | 0.45 | 0.18 | 0.02 | 0.96 | 0.95 |
| *HERC2* | 10:6217628 | A/G | snp | synonymous | 2056R | 44.9 | 0.97 | 0.87 | 0.50 | 0.30 | 0.95 | 0.85 | 0.50 | 0.21 | 1 | 0.96 |
| *HERC2* | 10:6216884 | G/A | snp | synonymous | 1999L | 4.3 | 0.95 | 0.92 | 0.70 | 0.47 | 1 | 0.89 | 0.57 | 0.33 | 1 | 0.97 |
| *HERC2* | 10:6218296 | C/T | snp | splice_region | NA | 4.3 | 0.87 | 0.83 | 0.32 | 0.04 | 0.92 | 0.63 | 0 | 0 | 0.92 | 0.98 |
| *KITLG* | 5:36821134 | TC/T | indel | 5′ UTR | NA | 3.1 | 0.03 | 0.03 | 0.05 | 0.03 | 0.06 | 0 | 0 | 0.02 | 0.20 | 0.40 |
| *MLANA* | 7:22505901 | A/T | snp | 3′ UTR | NA | 98.8 | 0.04 | 0.38 | 0.02 | 0.15 | 0 | 0.45 | 0 | 0.82 | 0 | 0 |
| *MLANA* | 7:22504421 | G/A | snp | splice_region | NA | 98.8 | 0.07 | 0.32 | 0.01 | 0.14 | 0 | 0 | 0 | 0.83 | 0 | 0 |
| *MLANA* | 7:22499952 | C/A | snp | missense | 18H>18N* | 98.8 | 0.04 | 0.30 | 0.04 | 0.10 | 0.06 | 0.02 | 0 | 0.84 | 0 | 0 |
| *MLANA* | 7:22505418 | T/C | snp | 3′ UTR | NA | 98.8 | 0.08 | 0.03 | 0.01 | 0.02 | 0 | 0 | 0 | 0.60 | 0 | 0.01 |
| *OCA2* | 10:6277903 | T/C | snp | missense | 126M>126T* | 47.9 | 0.34 | 0.55 | 0.10 | 0.01 | 0.59 | 0.66 | 0.24 | 0.09 | 0.73 | 0.08 |
| *OCA2* | 10:6231633 | G/A | snp | missense | 8R>8K* | 7.7 | 0.19 | 0.14 | 0.52 | 0.69 | 0.04 | 0.05 | 0.83 | 0.87 | 0 | 0.15 |

| | | | | | | | | | | | | | | | | |
|---|---|---|---|---|---|---|---|---|---|---|---|---|---|---|---|---|
| *OCA2* | 10:6231647 | C/T | snp | splice_region | NA | 7.3 | 0.81 | 0.81 | 0.22 | 0.05 | 0.81 | 0.90 | 0 | 0 | 1 | 0.83 |
| *OCA2* | 10:6234970 | T/C | snp | 5′ UTR | NA | 16.3 | 1 | 0.95 | 0.43 | 0.29 | 0.93 | 0.88 | 0.23 | 0.39 | 1 | 0.87 |
| *OCA2* | 10:6234919 | C/T | snp | 5′ UTR | NA | 16.3 | 0.99 | 0.87 | 0.19 | 0.05 | 0.79 | 0.87 | 0 | 0 | 1 | 0.87 |
| *SLC45A2* | 7:5253464 | G/C | snp | 3′ UTR | NA | 29.4 | 0.51 | 0.63 | 0.20 | 0.46 | 0.36 | 0.26 | 0.55 | 0.65 | 0.71 | 0.26 |
| *SLC45A2* | 7:5267285 | C/T | snp | synonymous | 204Q | 13.2 | 0.55 | 0.29 | 0.92 | 0.42 | 0.58 | 0.70 | 0.08 | 0.06 | 1 | 0.77 |
| *SLC45A2* | 7:5261935 | T/C | snp | missense | 368N>368D* | 54.6 | 0 | 0.08 | 0.86 | 0.23 | 0.43 | 0.81 | 0 | 0 | 0.07 | 0.80 |
| *SLC45A2* | 7:5256916 | T/C | snp | missense | 517M>517V* | 54.6 | 1 | 0.51 | 0.94 | 0.39 | 0.64 | 0.80 | 0.25 | 0.16 | 1 | 0.81 |
| *TYRP1* | 7:25294016 | T/C | snp | 5′ UTR | NA | 0 | 0 | 0.03 | 0.36 | 0.11 | 0.19 | 0.92 | 0 | 0.09 | 0.21 | 0.02 |
| *TYRP1* | 7:25297310 | G/T | snp | synonymous | 23A | 0.4 | 0.79 | 0.71 | 0.70 | 0.66 | 0.39 | 0.96 | 0.88 | 0.53 | 1 | 0.07 |
| *TYRP1* | 7:25305183 | T/A | snp | 3′ UTR | NA | 0.4 | 0.11 | 0.22 | 0.41 | 0.38 | 0.12 | 0.85 | 0.60 | 0.15 | 0 | 0.10 |
| *TYRP1* | 7:25304956 | A/G | snp | 3′ UTR | NA | 0.4 | 0.14 | 0.31 | 0.26 | 0.28 | 0.04 | 0.92 | 0.41 | 0.26 | 0.11 | 0.16 |
| *TYRP1* | 7:25305295 | C/T | snp | 3′ UTR | NA | 0.4 | 0.33 | 0.44 | 0.35 | 0.49 | 0.09 | 0.95 | 0.45 | 0.59 | 0.42 | 0.19 |
| *TYRP1* | 7:25300749 | T/C | snp | synonymous | 226H | 19.8 | 0 | 0.06 | 0 | 0.15 | 0.55 | 0.92 | 0 | 0.03 | 0 | 0.49 |
| *TYRP1* | 7:25305310 | C/A | snp | 3′ UTR | NA | 0.4 | 0 | 0 | 0.01 | 0.01 | 0.85 | 0 | 0.04 | 0 | 0 | 0.68 |
| *TYRP1* | 7:25305553 | T/C | snp | 3′ UTR | NA | 0.4 | 0 | 0 | 0 | 0.02 | 0.85 | 0 | 0 | 0 | 0 | 0.70 |
| *TYRP1* | 7:25305619 | A/C | snp | 3′ UTR | NA | 0.4 | 0 | 0 | 0 | 0.02 | 0.83 | 0 | 0 | 0.02 | 0 | 0.77 |
| *TYRP1* | 7:25305228 | A/G | snp | 3′ UTR | NA | 0.4 | 0 | 0.08 | 0 | 0.01 | 0.86 | 0 | 0 | 0 | 0 | 0.78 |
| *TYRP1* | 7:25305689 | T/C | snp | 3′ UTR | NA | 0.4 | 0.11 | 0.16 | 0 | 0.16 | 0.86 | 0 | 0.23 | 0.08 | 0.04 | 0.79 |
| *TYRP1* | 7:25306809 | G/A | snp | 3′ UTR | NA | 0.4 | 0 | 0.13 | 0 | 0.08 | 0.82 | 0 | 0.13 | 0.04 | 0.17 | 0.82 |
| *TYRP1* | 7:25304882 | T/C | snp | missense | 443S>443P* | 0.4 | 0.09 | 0.08 | 0.11 | 0.34 | 0.79 | 0 | 0.26 | 0.05 | 0.22 | 0.83 |
| *TYRP1* | 7:25306116 | T/A | snp | 3′ UTR | NA | 0.4 | 0.10 | 0.19 | 0 | 0.07 | 0.81 | 0 | 0.20 | 0 | 0 | 0.84 |
| *TYRP1* | 7:25306464 | C/A | snp | 3′ UTR | NA | 0.4 | 0.10 | 0.14 | 0.14 | 0.13 | 0.82 | 0 | 0.36 | 0.12 | 0 | 0.84 |
| *TYRP1* | 7:25306631 | C/G | snp | 3′ UTR | NA | 0.4 | 0.60 | 0.21 | 0.39 | 0.35 | 0.92 | 0 | 0.65 | 0.09 | 0.19 | 0.85 |

**Table S10. Differential transcript usage at candidate pigmentation genes.** For each candidate gene with at least two expressed isoforms, we tested DTU under four experimental designs: feather patch (43 samples), plumage color (males only, 34), sex (43), patch–color combination (42). Color groups and patch–color combinations represented by fewer than three samples were excluded. For each gene and test, the table shows the number of expressed isoforms, the adjusted *p*-value (Adjusted *p* gene; the smallest Benjamini–Hochberg adjusted *p*-value among the gene's isoforms) and the change, in percentage points, in the proportion of the dominant isoform between the groups in which it is most and least abundant. Isoform proportions for genes with differential transcript usage at adjusted $p < 0.001$ are shown in Fig. S4.

| Gene | Test | N isoforms | Adjusted *p* (gene) | Δ dominant (pp) | Dominant lowest in | Dominant highest in |
|---|---|---|---|---|---|---|
| *ASIP* | Patch | 8 | 0.16 | 22.3 | BR | CR |
| *GPT2* | Patch | 3 | 1.0 | 0.1 | BR | BA |
| *HERC2* | Patch | 5 | 0.26 | 2.5 | BA | CR |
| *KITLG* | Patch | 4 | 0.16 | 5.5 | BA | BR |
| *MLANA* | Patch | 2 | 0.005 | 0.5 | BR | CR |
| *OCA2* | Patch | 6 | 0.1 | 6 | CR | BA |
| *SLC45A2* | Patch | 4 | 0.012 | 12.8 | CR | BA |
| *TYRP1* | Patch | 8 | 0.052 | 12 | CR | BR |
| *ASIP** | Color (males) | 8 | 3.62e-5 | 52.4 | GR | SA |
| *GPT2* | Color (males) | 3 | 0.085 | 0.8 | SA | OR |
| *HERC2* | Color (males) | 5 | 0.88 | 2.6 | BC | OR |
| *KITLG* | Color (males) | 4 | 0.064 | 20 | BC | OR |
| *MLANA* | Color (males) | 2 | 0.0027 | 0.6 | SA | BC |
| *OCA2* | Color (males) | 6 | 0.085 | 10 | OR | BC |
| *SLC45A2* | Color (males) | 4 | 0.13 | 8.7 | BC | SA |
| *TYRP1* | Color (males) | 8 | 0.049 | 15.7 | BC | OR |
| *ASIP* | Sex | 8 | 0.41 | 14.9 | female | male |
| *GPT2* | Sex | 3 | 0.48 | 0.1 | male | female |
| *HERC2* | Sex | 5 | 0.48 | 0.7 | female | male |
| *KITLG* | Sex | 4 | 0.41 | 7.2 | male | female |
| *MLANA* | Sex | 2 | 0.015 | 0.4 | male | female |
| *OCA2* | Sex | 6 | 0.088 | 2.9 | male | female |
| *SLC45A2* | Sex | 4 | 0.21 | 6.6 | female | male |
| *TYRP1* | Sex | 8 | 0.088 | 3.5 | male | female |
| *ASIP** | Patch/color combination | 8 | 1.12e-5 | 84.5 | CR_BR | CR_BC |
| *GPT2* | Patch/color combination | 3 | 0.14 | 1 | BR_SA | BR_OR |
| *HERC2* | Patch/color combination | 5 | 0.46 | 5.1 | BR_BR | BR_SA |
| *KITLG* | Patch/color combination | 4 | 0.093 | 20.6 | BR_BC | BR_OR |
| *MLANA** | Patch/color combination | 2 | 5.81e-5 | 1.1 | BR_SA | CR_BR |
| *OCA2* | Patch/color combination | 6 | 0.037 | 16.8 | CR_GR | BR_BC |

| | | | | | | |
|---|---|---|---|---|---|---|
| *SLC45A2* | Patch/color combination | 4 | 0.036 | 17 | CR_BC | BA_SA |
| *TYRP1* | Patch/color combination | 8 | 0.052 | 26 | CR_BC | BR_OR |
